# eSPLIT and iSWAP: CRISPR-Mediated Conditional Allele Engineering with Short Artificial Introns

**DOI:** 10.64898/2025.12.22.696077

**Authors:** Ankeeta Koirala, Kyle O’Connor, Samantha Bare, Max Toubin, Anastasiya Vydra, Annelise Cassidy, Amalee Strupp, Destinee Thomas, Hanying Chen, Stephane Pelletier

## Abstract

Engineering conditional alleles remains a major challenge in functional genomics. Short Artificial Introns (SAIs) have emerged as powerful tools to simplify allele design and characterization, yet practical guidelines for their implementation remain limited. Here, we describe two streamlined strategies, eSPLIT (exon split) and iSWAP (intron swap), that enable efficient generation of conditional alleles using SAIs. In eSPLIT, a compact cassette containing essential intronic elements flanked by loxP sites is integrated within an exon, whereas in iSWAP, a native intron is replaced with an SAI. In the absence of Cre recombinase, the SAI is recognized as an intron and removed by the splicing machinery, allowing normal gene expression. Following Cre-mediated recombination, excision of critical intronic sequences disrupts splicing, leaving residual sequences, including stop codons in all reading frames, thereby causing premature translation termination and gene inactivation. We optimized these approaches by generating a series 18 *SCYL1* alleles in human cells, validated their general applicability in 5 mouse models across multiple genes and further extended the approach to the FLP-FRT system in vivo. By defining practical rules for SAI placement, we establish a robust and scalable framework for engineering conditional alleles, with broad utility in functional genomics and disease modeling.

**SHORT 100 – WORD ABSTRACT:** We introduce eSPLIT and iSWAP, two streamlined strategies for generating conditional alleles using a short artificial intron (SAI). eSPLIT integrates a loxP-flanked SAI within an exon, while iSWAP replaces a native intron. Cre-mediated excision disrupts SAI splicing, leading to gene inactivation. Optimized using *SCYL1* alleles in human cells and validated in mouse models across multiple genes, these strategies provide a simple and efficient approach for engineering conditional alleles. By outlining design rules for SAI placement, we provide a robust and scalable strategy for functional genomics and disease modeling.

## MAIN

Conditional alleles are indispensable tools in functional genomics, enabling spatial and temporal control of gene inactivation in model organisms. They allow researchers to bypass embryonic lethality, investigate gene function in specific tissues, cell types or developmental stages, and dissect complex biological processes. The most widely used method for generating conditional alleles relies on the Cre-loxP system, in which critical exons of a gene are flanked by loxP sites (“floxed”) and excised upon Cre recombinase expression. Despite its utility, the process of engineering such alleles is often labor-intensive, time-consuming, and technically challenging [3].

To overcome this limitation, we and others explored the use of short artificial introns (SAIs) as a streamlined approach for generating conditional alleles [4–7]. This technology enables conditional gene inactivation by facilitating a switch from an active to an inactive state [4–7]. Built upon the Cre-loxP system, current SAI technologies involve the insertion of a compact DNA cassette directly within an exon of the target gene. This cassette encodes a splice donor site, followed by essential intronic elements (including a branch point and a polypyrimidine tract) flanked by two loxP sites, and a splice acceptor site (**Fig. 1a**). In the absence of Cre recombinase, the cassette is recognized as an intron during RNA processing and is efficiently spliced out, allowing the production of a functional protein. Upon Cre-mediated recombination, intronic elements are excised, leaving behind residual sequences that prevent normal splicing, leading to gene inactivation (**Fig. 1a**). These synthetic introns are engineered to recapitulate the essential features of natural introns while minimizing sequence length. Cassette miniaturization, and the need for a single insertion event, as opposed to two for conventional designs, enhance the efficiency of allele generation and facilitate downstream characterization [4–7].

**Figure 1.**
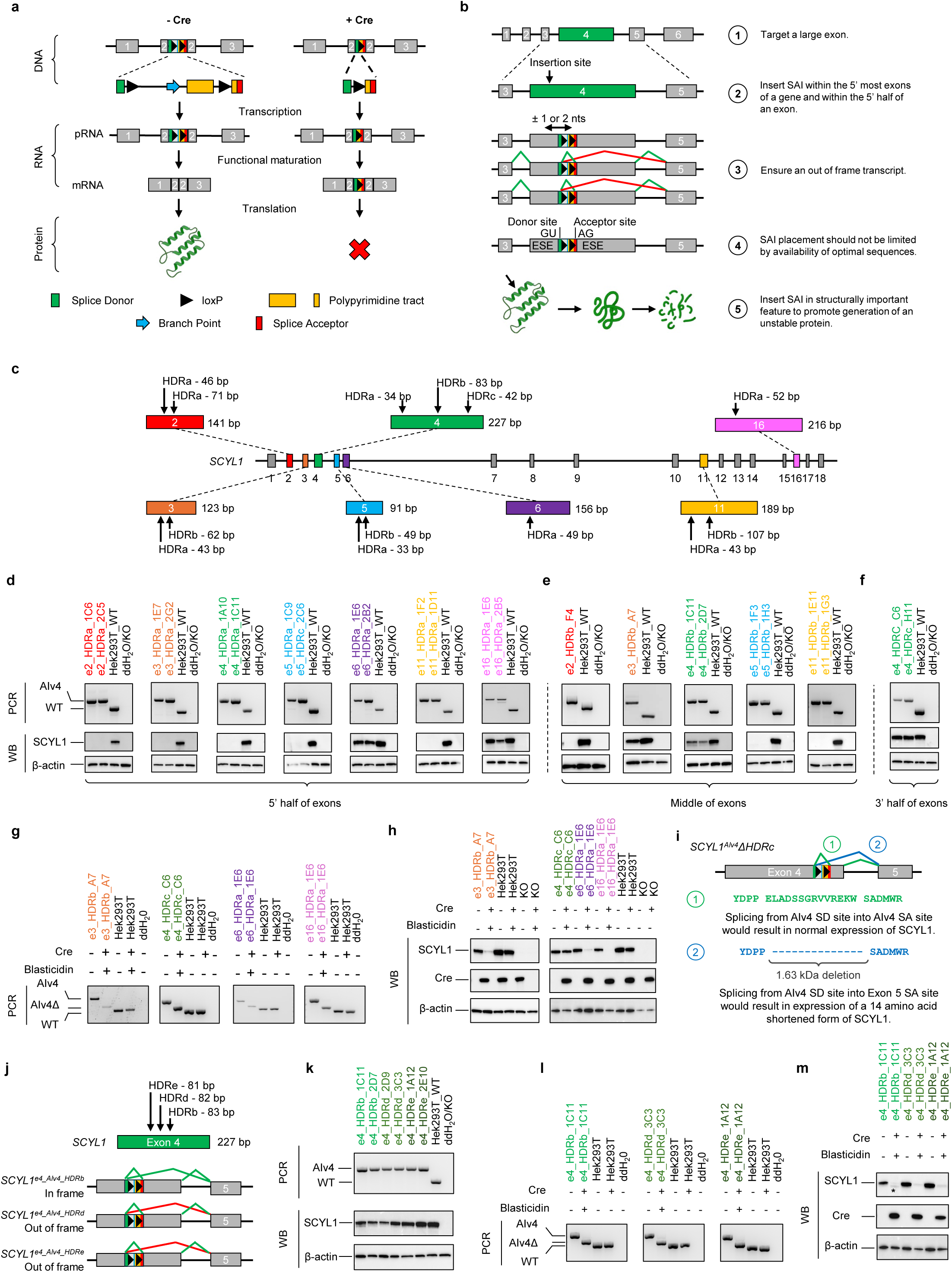
Optimization of SAI insertion sites for generating conditional alleles using the eSPLIT design. a) Schematic and principled of the eSPLIT strategy for creating conditional alleles in cell lines. The eSPLIT design, adapted from the DECAI approach, involves inserting a short cassette containing a splice donor, essential intronic sequences flanked by loxP sites, and a splice acceptor. In the absence of Cre, the cassette (Artificial Intron version 4; AIv4) is recognized as an intron and removed by the splicing machinery during transcript maturation, allowing normal gene expression. Upon Cre-mediated recombination between the loxP sites, the intron is disrupted and no longer recognized by the splicing machinery. Consequently, the retained intronic sequences, which harbor stop codons in all three reading frames, remain in the transcript and cause premature translation termination. b) Design considerations to be empirically tested. Previous design considerations stipulated that the AIv4 cassette should be inserted within the 5’ most exons (green boxes) of a gene and within the 5’ half of an exon (exon 4, highlighted in green) to promote mRNA degradation via the nonsense-mediated mRNA decay pathway (Popp and Maquat, 2016) (1 and 2). The artificial intron should be inserted such that potential splicing events driven by the AIv4 splice donor with downstream exon(s) results in an out of frame transcript (red line) (3). The location of the AIv4 cassette should not depend on the availability of optimal sequences upstream of the splice donor and downstream of splice acceptor sites (4). When possible, short AI should be introduced within sequences encoding structurally important feature of the encoded protein such that alterations within that structure would result in the generation of unfolded protein. Grey boxes, exons; grey lines; introns; green boxes, target exons; green line, proper splicing events; red lines, undesirable in-frame splicing events. c) Schematic representation of the SCYL1 gene and AIv4 cassette insertion sites. In total, 12 homozygous cell line models were engineered using CRISPR-Cas9 technology. In most cases, two clones were identified and analyzed for SCYL1 expression. Insertion sites included the 5′ half of exons 2, 3, 4, 5, 6, 11, and 16; the middle of exons 2, 3, 4, 5, and 11; and the 3′ half of exon 4. d) Effects of SAI insertion near the 5′ end of SCYL1 exons on gene expression. HEK293T cells harboring various eSPLIT designs (illustrated in panel C) were genotyped by PCR (upper panels) and validated by Sanger sequencing (Additional Figure 1), followed by assessment of SCYL1 protein expression in the absence of Cre recombinase (lower panels). Insertion of the AIv4 cassette within the 5′ half of exons 2, 3, 4, 5, and 11 resulted in loss of SCYL1 expression. β-actin was used as a loading control. e) Effects of SAI insertion near the middle of SCYL1 exons on gene expression. HEK293T cells harboring the indicated eSPLIT designs (panel C) were assessed for SCYL1 expression in the absence of Cre recombinase. Insertion of the AIv4 cassette into the middle of exons 3 and 4 preserved SCYL1 expression, whereas insertion into exons 2, 5, and 11 resulted in loss of expression. f) Effect of SAI insertion at the 3′ end of SCYL1 exon 4 on gene expression. HEK293T cells with AIv4 cassette insertion at the 3′ end of exon 4 expressed a slightly shorter form of SCYL1. g) Cre-mediated recombination of the functional eSPLIT designs. HEK293T cells harboring various eSPLIT designs were infected with lentiviruses expressing Cre recombinase together with mCherry and a blasticidin resistance gene. Following blasticidin selection, clones were analyzed for Cre-mediated recombination of the different allele designs. A shift in PCR banding patterns indicated near-complete recombination between the loxP sites within the AIv4 cassette in the presence of Cre recombinase. This was confirmed by Sanger sequencing (Additional Figure 2). h) SCYL1 protein expression following Cre-mediated recombination of the AIv4 cassette. In most eSPLIT designs, recombination resulted in loss of SCYL1 expression. Notably, insertion of the AIv4 cassette at the 3′ end of exon 4 still produced a shorter version of SCYL1 even in the presence of Cre recombinase. Also note the presence of a faint, shorter version of SCYL1 in the e4_HDRb clone in response to Cre expression. This likely represents expression of a shorter form of SCYL1 resulting from the splicing of the splice donor into the splice acceptor site of exon 5, as discussed in our previous publications[4, 5]. Cre expression was confirmed by western blotting and β-actin was used as loading controls. i) The shorter form of SCYL1 likely arises from splicing between the splice donor (SD) of the AIv4 cassette and the splice acceptor (SA) of exon 5. This in-frame splicing event removes 14 amino acids, producing a protein with a mobility shift equivalent to 1.63 kDa. j) Schematic representation of SCYL1 exon 4 and AIv4 cassette insertion sites. k) All three frame insertion designs permitted normal SCYL1 expression compared to the parental line in the absence of Cre recombinase. Two clones for each insertion site were identified by PCR genotyping, validated by Sanger sequencing (Additional Figure 3), and assessed for SCYL1 protein expression by western blotting. β-actin was used as a loading control. l) Cre-mediated recombination of the AIv4 cassette in validated clones. HEK293T cells harboring the indicated eSPLIT designs were infected with lentiviruses expressing Cre recombinase together with mCherry and a blasticidin resistance gene. Following blasticidin selection, clones were analyzed for Cre-mediated recombination. A shift in PCR banding patterns indicated near-complete recombination between the loxP sites within the AIv4 cassette in the presence of Cre recombinase. Recombination was confirmed by Sanger sequencing (Additional Figure 3). m) SCYL1 protein expression following Cre-mediated recombination of the AIv4 cassette. Unlike in-frame insertions, no shorter SCYL1 isoform was observed in eSPLIT designs where the AIv4 cassette was inserted out of frame. Cre expression was confirmed by western blotting, and β-actin was used as a loading control. The shorter form of SCYL1 observed in e4_HDRb, in the presence of Cre recombinase likely arises from splicing between the splice donor (SD) of the AIv4 cassette and the splice acceptor (SA) of exon 5. This in-frame splicing event removes 48 amino acids, producing a protein with a mobility shift equivalent to approximately 5 kDa.

Although the technology is still in its early developmental stages, we and others have proposed design guidelines for engineering such alleles using short artificial introns (SAIs), drawing primarily on current knowledge of splicing regulation, translation termination, and nonsense-mediated mRNA decay (NMD) [4–6]. However, in the absence of systematic experimental validation, these largely theoretical guidelines leave the broader applicability of the technology open to debate. Moreover, the full potential of this approach should extend beyond exon splitting to include replacing endogenous introns with SAIs to achieve similar functional outcomes, thereby expanding the applicability of the technology to genes with diverse architecture.

In this study, we advance the development of SAI-based conditional allele engineering by experimentally defining robust and generalizable design principles. Using *SCYL1* as prototypic gene, we generated a series of conditional alleles in human cells employing two complementary strategies: the exon SPLIT (eSPLIT) strategy, where a coding exon is separated into two exons by a SAI, and the intron SWAP (iSWAP) strategy, where an endogenous intron is replaced with a SAI. Through systematic evaluation of these designs, we identified key parameters that govern allele functionality. We further extended these findings to mouse models, demonstrating that SAI-based conditional alleles using eSPLIT and iSWAP strategies can be efficiently generated in vivo and reliably inactivated upon Cre expression. Additionally, we adapted the technology to the FLP-FRT recombinase system, further broadening its utility.

Together, our results establish a validated framework for the rational design of conditional alleles using short artificial introns. This platform streamlines allele design, enabling a versatile and scalable approach to conditional allele engineering in cultured cells and in vivo models. We anticipate that the strategies and guidelines presented here will accelerate the adoption of SAI-based designs across a wide range of biomedical research applications.

## RESULTS

### eSPLIT Strategy: Design Principles

In our pilot study on the use of short artificial introns (SAIs) for the generation of conditional alleles in mice using the eSPLIT design strategy, we proposed a set of design guidelines informed by our understanding of splicing regulation and the nonsense-mediated mRNA decay (NMD) ([4, 5] and **Fig. 1b**). These theoretical guidelines included the following principles: 1) The cassette should be inserted within an exon large enough to accommodate the SAI. 2) The SAI cassette should be inserted into one of the 5′-most exons and within the 5′ half of that exon to promote transcript degradation via the NMD pathway following recombination; 3) The cassette should be positioned such that any splicing events involving the SAI splice donor and downstream exons produce out-of-frame transcripts, thereby ensuring gene inactivation; 4) The placement of the artificial intron should not be constrained by the presence of optimal intronic motifs upstream of the splice donor or downstream of the splice acceptor; and 5) When possible, short AI should be introduced within sequences encoding a structurally important feature of the encoded protein such that alterations within this structure would result in the generation of an unstable protein. Although these guidelines appeared reasonable, they had not been empirically evaluated[4, 5]. Furthermore, reports from other laboratories using the technology yielded inconsistent outcomes which prompted us to empirically define and validate optimized guidelines for the use of SAI cassettes in conditional allele design (personal communications).

To evaluate the functional constraints associated with SAI cassette insertion, we engineered a series of conditional alleles in HEK293T cells by targeting the SAI cassette (referred here as Artificial Intron version 4, AIv4 [5]) to various coding exons of the *SCYL1* gene (**Fig. 1c**). Seven exons spanning both the 5’ and 3’ regions of the gene, exons 2 (e2), 3 (e3), 4 (e4), 5 (e5), 6 (e6), 11 (e11), and 16 (e16), were selected for insertion (**Fig. 1c**). The AIv4 cassette was inserted into the 5’ half of each exon using CRISPR-Cas9 technology, as previously described and detailed in the Methods section. Successful cassette integration and homozygosity of the targeted alleles were confirmed by PCR genotyping (**Fig. 1d**) and Sanger sequencing (**Additional file 1b**). Engineering details for all alleles, including guide sequences, HDR templates, genotyping primers, PCR genotyping banding patterns, and engineering methodologies are presented in **Additional table 1** and **2,** and Methods section.

To assess functionality, SCYL1 protein expression was examined by western blotting using previously validated antibodies against SCYL1 [4, 5, 8]. This antibody recognizes the following epitope of SCYL1 protein, spanning amino acids 706 to 726 (SSVEPPPEGTRLASEYNWGGA, **Additional file 1b**)[8]. Unexpectedly, we found that cassette insertion was tolerated only in exons 6 and 16 whereas integration into exons 2, 3, 4, 5, and 11 abolished detectable protein expression (**Fig. 1d**, bottom panels). Notably, although all insertions were located within the 5’ half of their respective exons, several were positioned very close to the 5’ end (<50 nucleotides). This suggested that proximity to the exon start site may have impaired proper recognition of the cassette by the splicing machinery and consequently inactivated the allele. In line with this, previous studies using large intron-trapping technologies have reported that such cassettes should be placed at least 50 nucleotides downstream of the 5’ end of the exon to support proper splicing[9].

To test whether inserting the cassette close to the 5’ end of an exon is detrimental, we generated an additional set of alleles in which the cassette was inserted closer to the center of various exons (e2, e3, e4, e5, and e11). This strategy restored SCYL1 protein expression in some cases, notably for exons 3 and 4 (**Fig. 1e**). However, exons 2, 5 and 11 remained incompatible with cassette integration.

To further evaluate the positional constraints of SAI cassette insertion, we tested whether targeting the AIv4 cassette near the 3’ end of an exon would support the generation of a functional *SCYL1* allele. The cassette was inserted within the 3’ half of exon 4 (**Fig. 1f**), which was selected because a prior insertion near the center of this exon had been functionally tolerated (**Fig. 1e**). Correct targeting was confirmed by PCR genotyping and Sanger sequencing (**Fig. 1f** and **Additional file 1**), and SCYL1 protein expression was analyzed by western blotting (**Fig. 1f**, bottom panel). Although SCYL1 protein was detected at levels comparable to the wild-type allele, suggesting a fully functional allele, a slight reduction in molecular weight was observed. This suggested a change in protein sequence. We hypothesize that insertion of the cassette near the 3’ end of the exon may have promoted aberrant splicing between the donor site of the artificial intron and the acceptor site of the downstream exon. In this case, the splicing event would remain in-frame but lead to the exclusion of 14 amino acids from the SCYL1 protein, consistent with the altered mobility observed by western blot (**Fig. 1i**, and below for additional details). These findings suggest that, as with insertions near the 5′ end, positioning the cassette too close to the 3′ end of an exon may be suboptimal.

To assess recombinase-responsiveness, we next tested whether Cre-mediated excision of the essential intronic sequence would result in SCYL1 inactivation. Clones harboring functional conditional alleles were infected with a Cre-expressing lentivirus as described in the Methods section and illustrated in **Additional file 2a** and **b**. Efficient recombination of the alleles was confirmed by PCR amplification of the loci (**Fig. 1g**) and Sanger sequencing (**Additional file 2c**) and corresponded with high transduction efficiency of the lentivirus (**Additional file 2b**). SCYL1 expression was assessed by western blotting (**Fig. 1h** and **1m** for e4_HDRb) and, as expected, Cre-mediated recombination of the AIv4 cassette led to gene inactivation in all but one configuration, confirming proper functioning of the conditional alleles. An exception to this was observed for the allele with cassette insertion near the 3’ end of the exon (e4_HDRc), where SCYL1 expression was not abolished following recombination. This is likely due to the in-frame splicing event described above (**Fig. 1h**) between the splice donor site of the cassette and the acceptor site of exon 5 as illustrated in **Fig. 1i**. Aberrant splicing between the SD site of the cassette into the SA site of exon 5 is in frame, likely preserving a functional transcript despite recombination.

Collectively, these findings demonstrate that the position of cassette insertion is a critical determinant for the successful generation of functional conditional alleles using the SAI strategy. Based on our results, we recommend targeting insertions within relatively large exons, positioning the cassette at least 50 nucleotides downstream of the 5’ end and at least 50 nucleotides upstream of the 3’ end. This spacing minimizes the risk of aberrant splicing and helps ensure effective gene inactivation following recombination.

Previous studies[4, 5] and data presented above demonstrated that insertion of the SAI cassette can give rise to aberrant splicing events originating from within the cassette itself. When such events are in-frame, they may produce truncated transcripts encoding shortened forms of the target protein. In these cases, cassette insertion may not result in a complete loss of function, but rather in a partial loss of function or the acquisition of aberrant protein properties. To determine whether enforcing an out-of-frame splicing event with downstream exons enhances allele inactivation, we engineered additional *SCYL1* alleles by inserting the SAI cassette in all three reading frames (**Fig. 1j**). Correct targeting of each allele was confirmed by PCR genotyping and Sanger sequencing (**Fig. 1k** and **Additional file 3a**). Western blot analysis showed that, prior to recombination, all three alleles expressed SCYL1 protein at levels comparable to the wild-type allele (**Fig. 1k** bottom panel). Following Cre-mediated recombination, we observed near-complete recombination of all alleles (**Fig. 1l**) and loss of SCYL1 protein expression in all cases (**Fig. 1m**). Of note, a small band of lower molecular weight (noted with an asterisk (*)) was also observed for the allele with an in-frame cassette insertion (e4_HDRb, **Fig. 1m**), consistent with expression of a truncated protein product. As previously reported in our *Scyl1^AIv4^* mouse model, the faint expression is likely due to protein instability as removing amino acids in this region of the protein result in an unstable protein[4, 5]. These findings support the conclusion that designing SAI cassette insertions to promote out-of-frame splicing relative to downstream exons is a more effective strategy for achieving functional gene inactivation in conditional allele design using the eSPLIT system. This also highlights the importance of targeting regions of proteins which if disrupted will also result in protein instability as previously suggested in our pilot study[4, 5].

Collectively, these findings demonstrate that the position of cassette insertion is a critical determinant for the successful generation of functional conditional alleles using the SAI strategy. Based on our results, we recommend targeting insertions within relatively large exons (the largest of the gene, regardless of its location within the gene), positioning the cassette at least 50 nucleotides downstream of the 5’ end and at least 50 nucleotides upstream of the 3’ end. This spacing minimizes the risk of aberrant splicing and helps ensure effective gene inactivation following recombination. Moreover, the cassette should be positioned such that potential splicing events between the splice donor of the cassette and subsequent exons of the gene are out of frame (see **Fig. 6** for a summary of design guidelines).

### eSPLIT in Mouse Model Engineering

Given that one of the primary goals of this technology is to streamline and accelerate the generation of mouse models with conditional alleles, we next sought to validate the eSPLIT design strategy in vivo. To this end, we engineered two mouse models utilizing the eSPLIT approach. The first model was designed using a suboptimal configuration, with the AIv4 cassette inserted 17 nucleotides from the 5’ end of exon 2 of the *Rufy1* gene (*Rufy1^AIv4-1^*, **Fig. 2a** and **2b**). In contrast, the second model was generated according to our empirically validated design guidelines, with the AIv4 cassette inserted 87 nucleotides from the 5’ end of exon 2 (*Rufy1^AIv4-2^*, **Fig. 2c**), a configuration predicted to preserve proper gene function. Both models were generated using CRISPR-Cas9 mediated genome editing, as described in the Methods section. Proper allele targeting was confirmed by PCR genotyping and Sanger sequencing (**Fig. 2b** and **Additional file 4**). As feared, mice harboring the suboptimal allele failed to express RUFY1 (*Rufy1^AIv4-1/ AIv4-1^*, **Fig. 2f**), phenocopying *Rufy1-*deficient mice (*Rufy1^Del76/Del76,^* **Fig. 2f**, right panel). In contrast, mice carrying the optimally designed allele expressed RUFY1 at levels comparable to wild-type controls (**Fig. 2f**). To determine whether Cre-mediated recombination of the optimally designed allele (*Rufy1^AIv4-2^*) resulted in gene inactivation, we crossed these mice to a Cre driver line (B6.C-Tg(CMV-cre)1Cgn/J) which express Cre under the control of the CMV promoter[10]. As shown in **Fig. 2g** and **2h**, homozygous recombined animals (*Rufy1^AIv4-2Δ/AIv4-2Δ^*, **Fig. 2g**, and **Additional file 4**) no longer express Rufy1 (**Fig. 2h**). Together, these results demonstrate that conditional allele design using the eSPLIT strategy is more likely to succeed when our design principles are applied.

**Figure 2.**
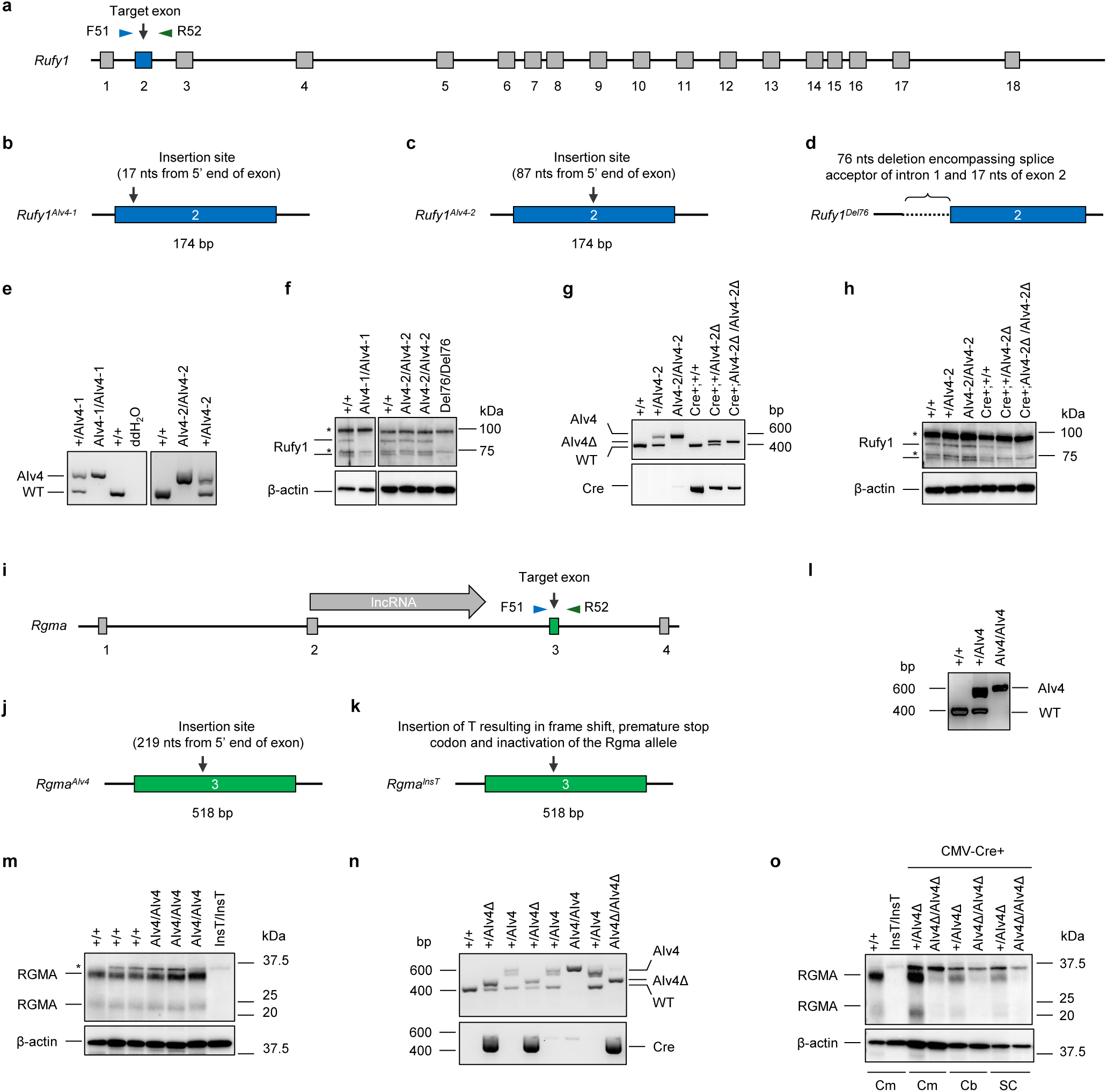
Generation of Rufy1^AIv4^ and *Rgma*^AIv4^ mouse models using eSPLIT design strategies. a) Schematic representation of *Rufy1* gene and AIv4 cassette insertion sites in exon 2. b) Schematic representation of *Rufy1*^AIv4-1^ allele design. AIv4 cassette was inserted 17 nucleotides from 5’ end of exon 2. c) Schematic representation of improved *Rufy1*^AIv4-2^ allele design. AIv4 cassette was inserted 87 nucleotides from 5’ end of exon 2. d) Schematic representation of *Rufy1*^Del76^ allele. A 76-nt deletion spanning intron 1-exon 2 of Rufy1 gene yielded a null allele. e) PCR genotyping of both *Rufy1^AIv4^* mouse models (*Rufy1^AIv4-1^* and *Rufy1^AIv4-2^*) illustrating correct allele targeting, validated by Sanger sequencing (Additional Figure 4). f) Western blot analysis of RUFY1 protein expression in wild-type mice, homozygous mice for both eSPLIT designs (*Rufy1^AIv4-1/AIv4-1^* and *Rufy1^AIv4-2/AIv4-2^*), and *Rufy1*-deficient mice (*Rufy1^Del76/Del76^*). The Rufy1 antibody recognizes two bands of approximately 90 kDa and 70 kDa, absent in *Rufy1^Del76/Del76^* cerebrum lysates. These correspond to the known isoforms RABIP4 and RABIP4′ of RUFY1 which differ by an N-terminal extension of 108 amino acids in RABIP4′. Two additional nonspecific bands (indicated with asterisks (*)) of ∼100 kDa and ∼75 kDa were also detected. Notably, mice homozygous for the suboptimal eSPLIT design (*Rufy1^AIv4-1/AIv4-1^*) lacked both RUFY1 isoforms (left panel), whereas homozygous mice for the optimal eSPLIT design (*Rufy1^AIv4-2/AIv4-2^*) expressed both isoforms. g) Cre-mediated recombination of the AIv4 cassette. *Rufy1^AIv4-2^* mice were crossed with CMV-Cre+ mice which express Cre ubiquitously, and recombination was detected as a shift in PCR banding pattern corresponding to excision between the loxP sites, which was confirmed by Sanger Sequencing (**Additional Fig. 4**). h) RUFY1 protein expression in wild-type, heterozygous, and *Rufy1^AIv4-2/AIv4-2^* mice, as well as their CMV-Cre-expressing counterparts. Both RUFY1 isoforms were absent in cerebrum lysates from CMV-Cre+; *Rufy1^AIv4-2Δ/AIv4-2Δ^* mice. β-actin was used as a loading control. i) Schematic representation of Rgma gene and AIv4 cassette insertion sites in exon 3. j) Schematic representation of *Rgma*^AIv4^ allele design. AIv4 cassette was inserted 219 nucleotides from 5’ end of exon 3. k) Schematic representation of *Rgma^InsT^* allele. A single-T insertion in exon 3 introduced a frameshift and premature stop, yielding a protein-null allele. l) PCR genotyping of Rgma mouse model (*Rgma^AIv4^*) illustrating correct allele targeting, validated by Sanger sequencing (**Additional Figure 4**). m) Western blot analysis of Rgma protein expression in wild-type mice, homozygous mice for *Rgma^AIv4/AIv4^*, and *Rgma*-deficient mice (*Rgma^InsT/InsT^*). The Rgma antibody recognizes two bands of approximately 35 kDa and 22 kDa, absent in *Rgma^InsT/InsT^* cerebrum lysates. These correspond to the two known isoforms of RGMA. One additional nonspecific band (∼37 kDa) was also detected. Notably, homozygous *Rgma^AIv4/AIv4^* mice express both Rgma isoforms. n) Cre-mediated recombination of the AIv4 cassette. *Rgma^AIv4^* mice were crossed with CMV-Cre+ mice, and recombination (*Rgma^AIv4Δ/AIv4Δ^)* was detected as a shift in PCR banding pattern corresponding to recombination between the loxP sites, which was confirmed by Sanger Sequencing (Additional Fig. 4h). o) RGMA protein expression in wild-type, Rgma-deficient (Rgma*^InsT/InsT^*) as well as CMV+ heterozygous, or CMV-Cre+;*Rgma^AIv4/AIv4^* mice. Both RGMA isoforms were absent in Cerebrum,(Cm) Cerebellum (Cb) or Spinal cord lysates from CMV-Cre+;*Rgma^AIv4Δ/AIv4Δ^* mice. β-actin was used as a loading control.

To further validate the robustness of this approach, we engineered a conditional allele of *Rgma* following our functionally validated eSPLIT design rules. As shown in **Fig. 2i** and **2j**, the AIv4 cassette was inserted 219 nucleotides downstream of the 5’ end and 299 nucleotides upstream of the 3’ end of Exon 3. Exon 3 was chosen as site of insertion as it is one of the largest exons of the *Rgma* gene, and sequences encoding Exon 2 also encode part of a long noncoding RNA transcript (**Fig. 2i**). PCR genotyping and Sanger sequencing confirmed the proper insertion of the cassette (**Fig. 2l** and **Additional file 4**). Mice homozygous for this allele expressed RGMA at levels comparable to wild type animals (**Fig. 2m**). Upon Cre-mediated recombination, via crossing onto the CMV-Cre mice (B6.C-Tg(CMV-cre)1Cgn/J) and generation of homozygous recombined animals (*Rgma^AIv4Δ/AIv4Δ^*, **Fig. 2n**), RGMA expression was significantly reduced, confirming functional inactivation of the allele (**Fig. 2o**). Additional models were engineered using these guidelines and prove to function as anticipated (data not shown). Together, these findings support the applicability and effectiveness of our optimized eSPLIT design guidelines for engineering functional conditional alleles in vivo.

### iSWAP Strategy: Design Principles

The findings described above provide important design principles for generating conditional alleles using the SAI cassette by targeting exon sequences. While this approach is relatively straightforward to implement and has proven effective, its utility is limited by exon size and cassette positioning, as shown in the first part of this manuscript. To address this limitation, we developed an alternative strategy, termed intron SWAP (iSWAP), in which a native intron is replaced with the AIv4 cassette (**Fig. 3a**). Similar to eSPLIT, in the absence of Cre mediated recombination, the cassette is removed during functional maturation of the transcript, allowing for normal protein expression. In the presence of Cre, the cassette is rearranged, remains within the transcript and impairs protein translation (**Fig. 3b**). We hypothesized that this approach would preserve the endogenous gene function prior to recombination while enabling efficient and predictable gene inactivation upon Cre-mediated excision of essential intronic sequences.

**Figure 3.**
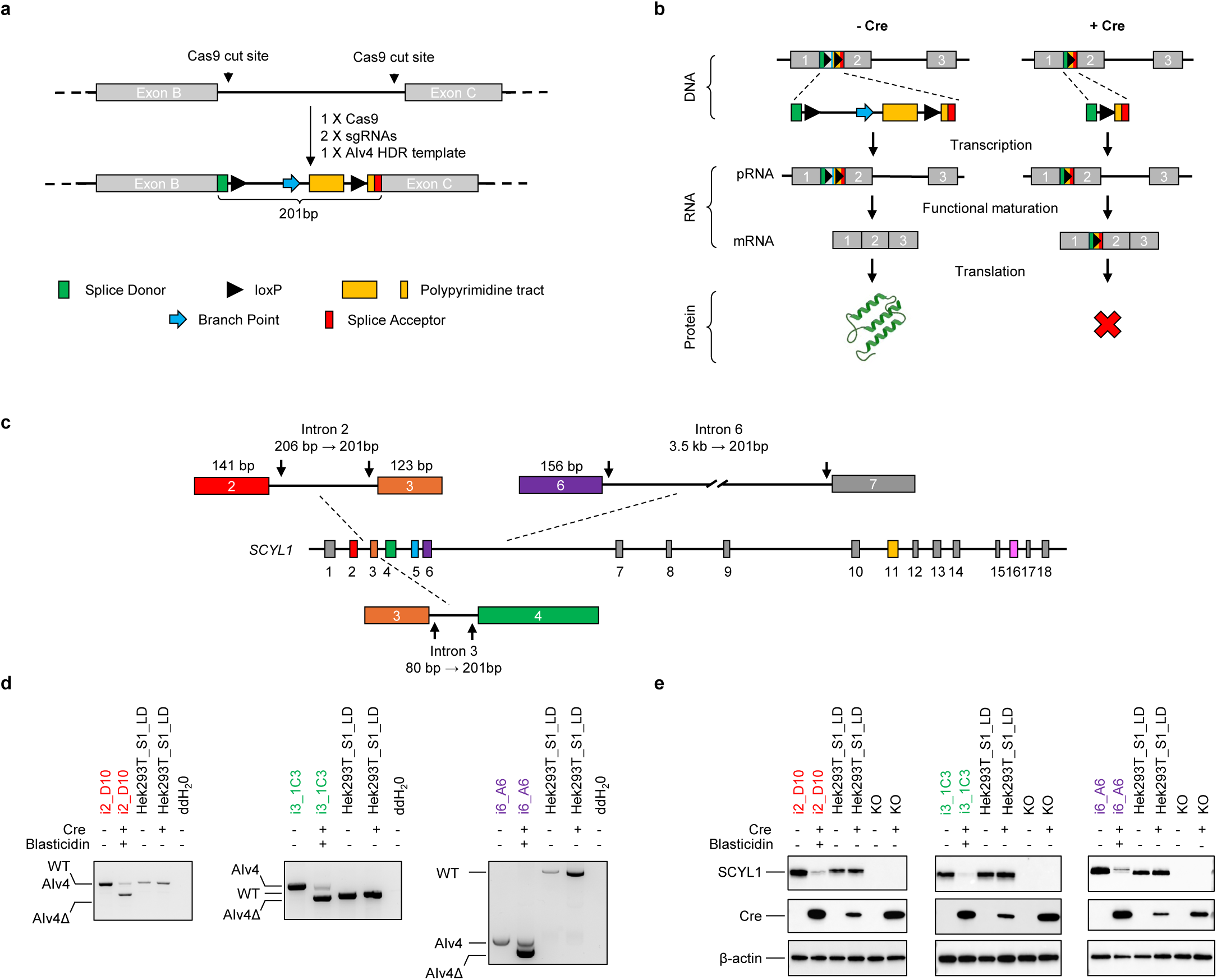
Generation of conditional alleles using the iSWAP design. a) Schematic of the iSWAP strategy for creating conditional alleles in cell lines. The iSWAP design, adapted from the DECAI approach, involves replacing an endogenous intron with a short cassette containing a splice donor, essential intronic sequences flanked by loxP sites, and a splice acceptor. The SAI cassette used for the iSWAP and eSPLIT designs is the same. b) Principles of the iSWAP strategy. In the absence of Cre, the cassette (Artificial Intron version 4; AIv4) is recognized as an intron and removed by the splicing machinery during transcript maturation, allowing normal gene expression. Upon Cre-mediated recombination between the loxP sites, the intron is disrupted and no longer recognized by the splicing machinery. Consequently, the retained intronic sequences, which harbor stop codons in all three reading frames, remain in the transcript and cause premature translation termination. c) Schematic representation of the *SCYL1* gene and endogenous introns 2, 3 or 6 replacements. The replacement of endogenous introns requires the use of two sgRNAs and 1 homology directed repair template containing sequences encoding the artificial intron and homology arms of approximately 50 nucleotides (**Additional Table 1**). d) Engineering and functionality of iSWAP designs. Cre-mediated recombination of functional. Cell line models were engineered using CRISPR-Cas9 technology. In this case, alleles were engineered Hek293T cells containing a single copy of the *SCYL1* gene (*SCYL1_LD*) for ease of engineering. Engineering of Hek293T *SCYL1^+/LD^* (*Hek293T_S1_LD*) cell line is described in Methods section and engineering data presented in **Additional Fig. 5**. PCR genotyping and Sanger Sequencing (**Additional Fig. 5**) confirmed the proper engineering of the alleles. HEK293T cells harboring various iSWAP designs were infected with lentiviruses expressing Cre (+) recombinase together with mCherry and a blasticidin resistance gene. Following blasticidin selection, clones were analyzed for Cre-mediated recombination of the different allele designs. A shift in PCR banding patterns indicated significant recombination between the loxP sites within the AIv4 cassette in the presence of Cre recombinase. This was confirmed by Sanger sequencing (**Additional Fig. 5**). e) SCYL1 protein expression following Cre-mediated recombination of the AIv4 cassette in iSWAP alleles. Cre-mediated recombination resulted in loss of SCYL1 expression in all iSWAP alleles, corresponding to allele recombination frequency in that experiment. Cre expression was confirmed by western blotting and β-actin was used as loading controls.

To evaluate the efficacy of the iSWAP strategy in vitro, we generated conditional *SCYL1* alleles in human cells by replacing introns 2 (i2), 3 (i3), or 6 (i6) with the AIv4 cassette (**Fig. 3c**). In this case, two guide RNAs were used to excise the native intron and a single HDR molecule was used to insert the SAI. Details about HDR template design and guide selection are provided in the Methods section. For the engineering of these cell line models, we took advantage of a developed HEK293T cell line in which a single *SCYL1* allele is present. Details about engineering of this cell line are also presented in **Suppl Fig. 5** and Methods section. The proper engineering of all cell line models were confirmed using PCR amplification of the locus and Sanger sequencing (**Fig. 3d** and **Additional file 5**). In all cases, SCYL1 expression was preserved in the absence of Cre recombinase (**Fig. 3e**), indicating that gene function was maintained prior to recombination. Following Cre-mediated recombination, SCYL1 gene recombination was near complete and correspondingly protein expression was efficiently abolished, confirming successful gene inactivation (**Fig. 3e**). These results validate iSWAP as an efficient strategy for engineering conditional alleles, expanding the applicability of SAIs for the engineering conditional alleles in cultured human cells.

To further extend the practicality of the approach to animal models, we engineered a mouse model bearing a conditional allele of *Nipsnap1* (**Fig. 4a**). This gene was selected due to the size of its exons, which vary from 32 nucleotides to 141 nucleotides, making it challenging to engineer conditional alleles using eSPLIT design strategy. This mouse model was engineered by replacing the native intron 2 (i2) of the *Nipsnap1* gene with the AIv4 cassette using CRISPR-Cas9 technology (**Fig. 4a**). The proper engineering of the allele was confirmed by PCR amplification of the locus and Sanger sequencing (**Fig. 4b** and **Additional file 6**). The proper functioning of the allele was assessed by evaluating NIPSNAP1 protein expression by western blotting. As shown in **Fig. 4c**, mice homozygous for the AIv4 allele expressed NIPSNAP1 to levels comparable to wild type animals. To test recombination efficiency, mice bearing the *Nipsnap1^i2AIv4^* allele (*Nipsnap1^i2AIv4/i2AIv4^*) were crossed to the CMV-Cre driver mouse model (B6.C-Tg(CMV-cre)1Cgn/J). Efficient recombination of the AIv4 cassette was confirmed using PCR amplification of the locus and Sanger sequencing (**Fig. 4d** and **Additional file 6**). Consistent with the proper functioning of the allele, we also found that NIPSNAP1 expression was drastically reduced in mice homozygous for the recombined allele (**Fig. 4c**), correlating with allele recombination frequency shown in **Fig. 4b**. Antibody specificity was confirmed using protein extracts from a Nipsnap1 knockout animal (*Nipsnap1^Del230/Del230^*), engineered as detailed in the Methods section and **Additional tables 1** through **5**. Together, these findings validate iSWAP as an effective strategy for conditional allele engineering, including for genes in which short exon lengths preclude the use of the eSPLIT strategy. Again, the single insertion event simplifies design strategy and likelihood of successfully engineering the allele compared to conventional design strategies.

**Figure 4.**
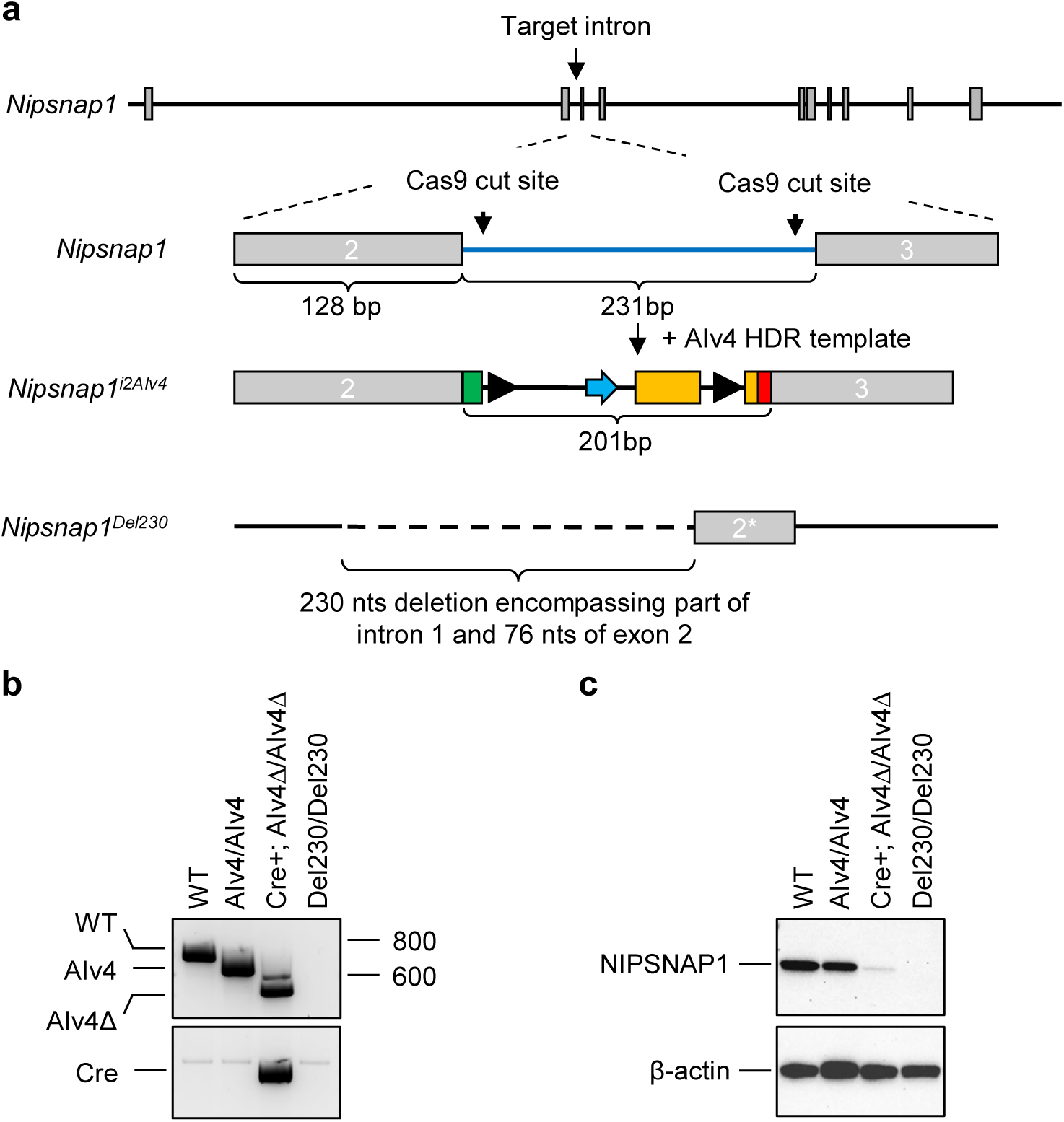
Generation of a mouse model bearing a conditional allele for Nipsnap1 using iSWAP design strategy. a) Schematic representation of the *Nipsnap1* gene and intron 2 replacement for the AIv4 cassette. b) PCR genotyping of *Nipsnap1^i2AIv4^* mouse model, illustrating correct allele targeting, validated by Sanger sequencing (**Additional Fig. 6b**). c) Western blot analysis of Nipsnap1 protein expression in wild-type mice, *Nipsnap1*^AIv4/AIv4^, Cre+ Nipsnap1AIv4//AIv4 and *Nipsnap1*-deficient mice (*Nipsnap1^Del230/Del230^*). Notably, mice homozygous *Nipsnap1^i2AIv4/i2AIv4^* express NIPSNAP1 to levels that are comparable to wild type.

### Extending SAI technology to the FLP/Frt system

To broaden the utility of this platform, we further adapted the SAI technology to the FLP-FRT recombinase system. We engineered a conditional allele of the *Stmn2* gene by inserting a modified SAI cassette termed Artificial Intron version 5 (AIv5), in which essential intronic elements are flanked by FRT sites as well as a splice donor and acceptor sites (**Fig. 5a**). This mouse model was engineered using CRISPR-Cas9 technology following eSPLIT design guidelines, the details of which are provided in the Methods section. The AIv5 cassette was inserted in exon 3 of the *Stmn2* gene, 81 nucleotides from the 5’ end of the exon and 92 nucleotides from its 3’ end (**Fig. 5b**). Proper engineering of the allele was confirmed by PCR amplification of the locus and Sanger sequencing (**Fig. 5c** and **Additional file 7**). Following the identification of multiple founders, some of them were outbred to wildtype mice for two generations. F2 mice were then crossed together to generate mice homozygous for the AIv5 allele. Western blot analyses from these mice revealed that STMN2 is expressed to levels comparable to those found in wild type animals (**Fig. 5d**). Crossing *Stmn2^AIv5^* mice to a FLP-deleter mouse line (B6.Cg-Tg(Pgk1-flpo)10Sykr/J[11]) resulted in the efficient recombination between the two Frt sites (**Fig. 5c** and **Additional file 7**) and abrogated STMN2 expression in homozygous mice (**Fig. 5d**). *Stmn2^LI/LI^* mice served as whole body knockout controls. The *Stmn2^LI/LI^* were engineered at the same time as the *Stmn2^AIv5^* model. Engineering details are provided in **Additional file 7** and Methods section. These findings expand the applicability of SAI designs to other recombinase systems. Future work will involve the generation of models using iSWAP with various recombinase systems, including Dre-Rox, Vika-Vox and others.

**Figure 5.**
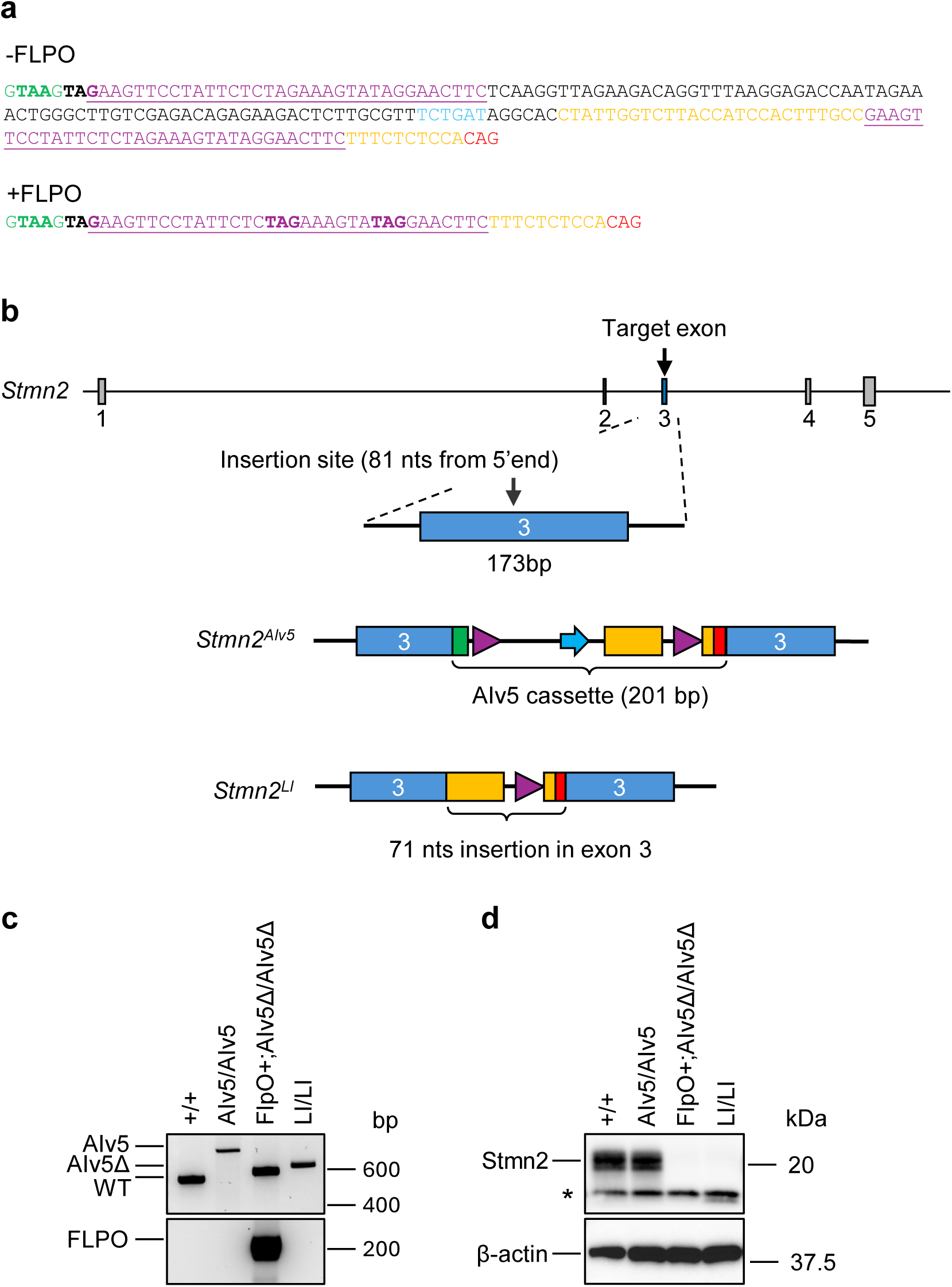
Extension of the SAI technology to the FLP/Frt system. a) Sequence of the unrecombined and recombined AIv5 cassette. Sequence of the AIv5 cassette, illustrating the substitution of the loxP sites from the AIv4 cassette for Frt sites. Of note, upon FLPO-mediated recombination, STOP codons in only 2 frames remains. This needs to be considered when engineering alleles using this design strategy. b) Schematic representation of the *Stmn2* gene, *Stmn2^AIv5^* and *Stmn2^LI^* alleles. c) PCR genotyping of wildtype, *Stmn2^AIv5/AIv5^*, FLPO+;*Stmn2^AIv5Δ/AIv5Δ^* and *Stmn2* deficient (*Stmn2^LI/LI^*) mice. Alleles were validated by Sanger sequencing (**Additional Fig. 7b and 7c**). d) Functional validation of the *Stmn2^AIv5^* allele. Western blot analysis of STMN2 protein expression in wild-type, *Stmn2^AIv5/AIv5^*, FLPO+;*Stmn2^AIv5Δ/AIv5Δ^*, and *Stmn2-*deficient mice (*Stmn2^LI/LI^*). *Stmn2^AIv5/AIv5^ mice* express STMN2 to levels that are comparable to wild-type mice whereas expression of STMN2 in FLPO+;*Stmn2^AIv5Δ/AIv5Δ^* mice is undetectable. The *Stmn2^LI/LI^* mouse model validated the STMN2 antibody.

Collectively, our findings define empirically validated design guidelines for the use of SAIs in generating conditional alleles in both cultured cells and in vivo. By introducing and systematically testing the eSPLIT and iSWAP strategies, we significantly expand the versatility and efficiency of SAI-based engineering of conditional alleles, providing multiple validated design approaches. Furthermore, we demonstrate that these strategies are compatible with multiple recombinase systems, thereby broadening their applicability. Guidelines for the design and engineering of these alleles are presented in **Fig. 6** and stipulate that: (1) The cassette should be inserted within an exon large enough to accommodate the AI; (2) The cassette does not have to be inserted within the 5’ most exons of a gene and within the 5’ half of an exon to inactivate the allele. Rather, the cassette should be inserted in the largest exon of a gene; (3) The artificial intron should be inserted such that potential splicing events driven by the SAI splice donor with downstream exon(s) results in an out of frame transcript; (4) The placement of the artificial intron should not be limited by the availability of optimal sequences upstream of the splice donor and downstream of splice acceptor sites as no correlation between allele functionality and ESEs could be drawn; (5) When possible, SAIs should be introduced within sequences encoding a structurally important feature of the encoded protein such that alterations within this structure would result in the generation of an unstable protein; and (6) Replacing endogenous introns with the SAI broadens the use of this technology for genes of various structures, including those with short exons.

**Figure 6.**
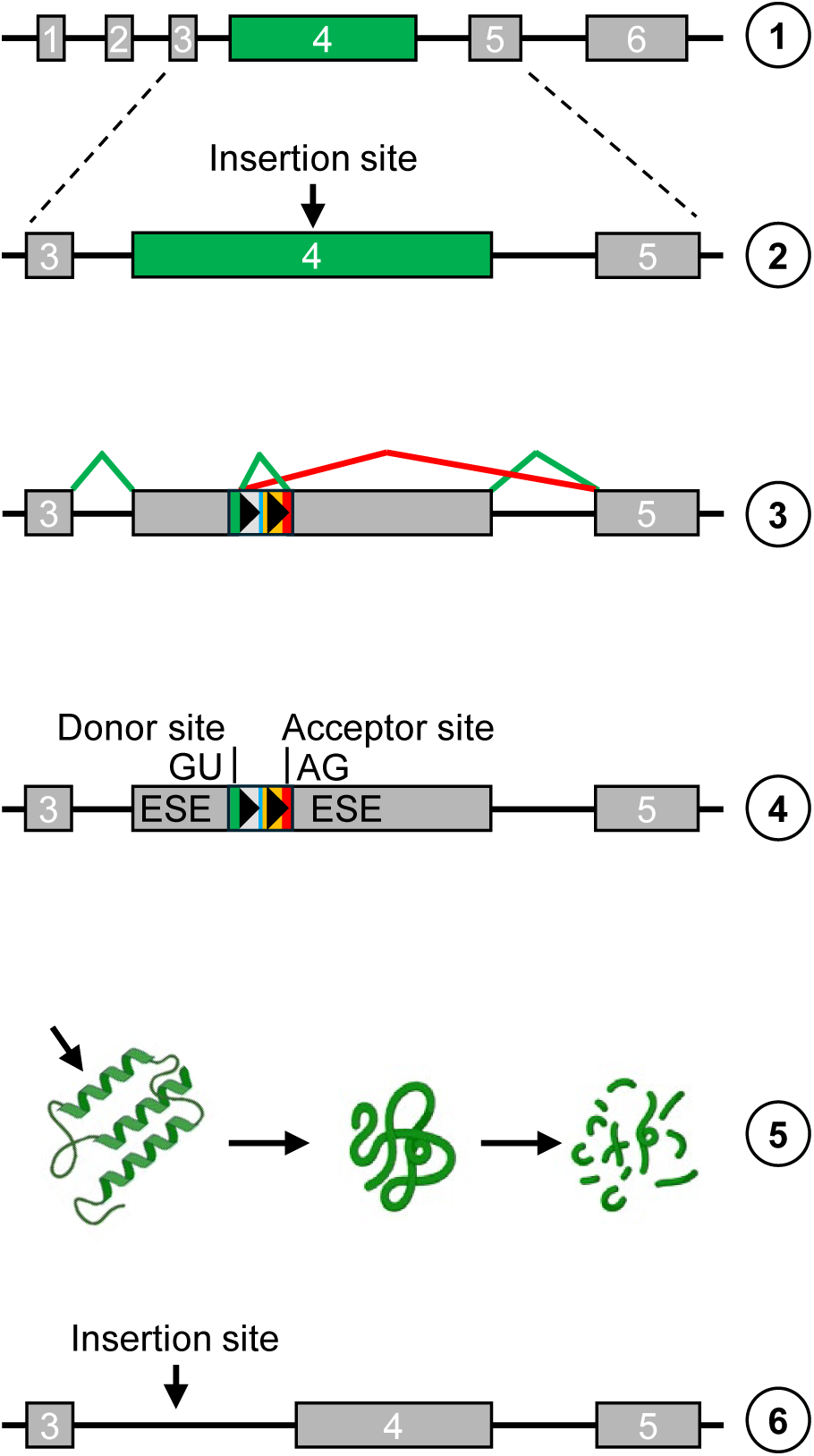
eSPLIT and iSWAP design guidelines. The SAI cassette should be inserted within an exon large enough to accommodate the artificial intron (AI). While placement within the 5′-most exons and the 5′ half of an exon can promote mRNA degradation via the nonsense-mediated decay (NMD) pathway, the cassette can be inserted into any exon - preferably those in the 5′ region - and should be positioned near the center of the exon. The artificial intron should be oriented so that any splicing driven by the AIv4 splice donor to downstream exons generates an out-of-frame transcript. The insertion site should not be constrained by the presence of canonical upstream or downstream splice site sequences; AIv4 functions reliably across a range of sequence contexts. When possible, the AI should be inserted into a coding region representing a structurally important domain of the encoded protein, such that perturbation would lead to an unstable or non-functional protein. The SAI cassette can replace any intron in the gene, but ideally one located near the 5′ end such that the nonsense mRNA decay pathway is engaged.

## DISCUSSION

The engineering of conditional alleles in mammalian systems remains a significant technical and financial challenge. Conventional approaches typically rely on flanking one or more critical exons with site-specific recombinase recognition sequences (SSRSs), such as loxP sites[12]. Upon expression of a site-specific recombinase (for example Cre, when using the Cre-loxP system), the targeted exons are excised, resulting in gene inactivation. Although this strategy has enabled numerous insights into gene function over the past four decades, the process of generating these alleles remains complex, time-consuming, and costly, making conditional alleles among the most difficult and expensive types to produce[12–14].

To overcome the limitations of traditional conditional allele engineering, artificial introns have been developed and incorporated into alternative strategies. Several iterations have emerged, including the COIN system, CRISPR-FLIP, and SAI systems[4–7, 9, 15]. The COIN approach employs an optimized conditional gene trap module (the COIN module) inserted in the antisense orientation relative to the target gene. Following Cre-mediated recombination, the module is inverted, leading to the expression of a reporter gene and premature termination of translation of the target transcript. Similarly, CRISPR-FLIP utilizes an inverted puromycin-STOP cassette positioned antisense to the target gene; Cre-mediated inversion results in allele inactivation. Both strategies are compatible with conventional and CRISPR-assisted gene targeting in embryonic stem (ES) cells and have been successfully used to generate numerous conditional mouse models. However, these methods remain inefficient, labor-intensive, and costly. Furthermore, the presence of strong promoters within reporter cassettes can influence the expression of neighboring genes, leading to unexpected phenotypes unrelated to the target gene[16].

As introduced earlier, SAI-based approaches aim to streamline the generation of conditional alleles in both animal models and cell lines. Despite their promise, the broader adoption of these technologies has been hindered by a lack of standardized design guidelines and limited systematic evaluation in both cultured cells and in vivo systems.

In this study, we provide a framework for the rational design and application of two related, SAI-based strategies: eSPLIT, which involves the insertion of a short artificial intron within an endogenous exon, as originally intended[7], and iSWAP, a novel strategy which involves the replacement of a native intron with an SAI. Our data demonstrate that these strategies are not only straightforward to design but also yield decent insertion efficiencies in both cultured cells and embryos. Importantly, allele characterization is markedly simplified, requiring only PCR amplification of the targeted locus followed by Sanger sequencing, as opposed to the extensive validation required for conventional conditional designs[12]. We further demonstrate that both eSPLIT and iSWAP alleles are functional and recombinase-responsive in cultured cells and in vivo. Our guidelines yield predictable outcomes, consistent with the intended design in the tested cases.

Each method has unique advantages. The eSPLIT strategy is broadly applicable, including for single-exon genes that are typically intolerant to conventional conditional strategies. However, its use is constrained by exon size and positioning of the cassette within an exon. In contrast, the iSWAP strategy is ideal for genes of various architectures, including those with short exons. Although iSWAP involves two genomic cuts, similar to the traditional strategy of flanking an exon with SSRSs[12], the recombination-mediated replacement of the intervening sequence with an SAI template is highly efficient. For example, we identified 5 correctly targeted founder animals out of 32 pups produced from a single round of electroporation for the engineering of the Nipsnap1 model.

Another notable advantage of both strategies is their compatibility with zygote electroporation. The short length of the HDR templates (∼300 nucleotides) facilitates efficient nuclear entry in pronuclear-stage zygotes, eliminating the need for microinjection. In contrast, longer single-stranded or double-stranded DNA templates often require microinjection or viral delivery due to their size and structural complexity. Moreover, both microinjection and viral delivery approaches present their own limitations[12]. It is unknown whether eSPLIT and iSWAP are compatible with iGONAD[17] but given the size of the HDR templates, both design strategies should be compatible.

In sum, the advantages of SAI-based conditional allele engineering include:

- Simplified design and validation – Alleles can be easily characterized using standard PCR and sequencing approaches.
- High editing efficiency – Both eSPLIT and iSWAP demonstrate robust targeting efficiency in cultured cells and embryos/mice.
- Reduced cost and time – Short HDR templates and electroporation-compatible designs eliminate the need for custom microinjection or viral vectors for delivery. A single session of electroporation and embryo manipulation is typically sufficient, reducing the number of fertilized eggs and mice used to produce mouse models with conditional alleles, adhering to the 3R principles.
- Broad applicability – eSPLIT enables conditional allele generation in genes that are poorly suited to traditional strategies, such as single-exon genes.
- Predictable outcomes – There is concordance between the intended design and in vivo outcomes, when following the proposed guidelines.
- Compatibility with multiple recombinases – The designs support flexibility in recombinase choice (e.g., Cre, Flp), enabling intersectional genetics and enhanced temporal/spatial control.

Together, these features make SAI-based strategies a compelling alternative or supplement to conventional methods for generating conditional alleles. By lowering technical and financial hurdles, they have the potential to greatly expand access to conditional genetics across diverse model systems and research laboratories.

Given that the goal of this manuscript is to define practical guidelines for the broader application of SAI strategies for conditional allele generation, we outline the following design principles for engineering eSPLIT and iSWAP alleles (**Fig. 6**):

eSPLIT Design Guidelines:

- The SAI cassette should be inserted within an exon large enough to accommodate the artificial intron (AI).
- While placement within the 5′-most exons and the 5′ half of an exon can promote mRNA degradation via the nonsense-mediated decay (NMD) pathway, the cassette can be inserted into any exon — preferably those in the 5′ region — and should be positioned near the center of the exon.
- The artificial intron should be oriented so that any splicing driven by the AIv4 splice donor to downstream exons generates an out-of-frame transcript.
- The insertion site should not be constrained by the presence of canonical upstream or downstream splice site sequences; AI functions reliably across a range of sequence contexts.
- When possible, the AI should be inserted into a coding region representing a structurally important domain of the encoded protein, such that perturbation would lead to an unstable or non-functional protein.

iSWAP Design Guidelines:

- The SAI cassette can replace any intron in the gene, but ideally one located near the 5′

end such that the nonsense mRNA decay pathway is more likely engaged.

In all cases, a thorough understanding of gene structure and regulatory elements is essential for the proper engineering of conditional alleles using these design strategies, as with any genome engineering projects. Disruption of regulatory elements or targeting known spliced exons should be avoided. Models should be rigorously validated at the genomic level and, critically, at the protein level to confirm functional inactivation.

In conclusion, the eSPLIT and iSWAP design strategies represent a significant advance in the generation of conditional alleles, offering a simplified, efficient, and cost-effective alternative to traditional methods. By leveraging compact, electroporation-compatible templates and flexible cassette designs, these approaches reduce both technical barriers and resource requirements.

## METHODS

### Antibodies, oligonucleotides, RNA transcripts and other reagents

All reagents used in this study are listed in **Additional Tables 1 through 7**.

### Tissue culture

Hek293T cells were obtained from ATCC (CRL-3216). Cells were maintained in Dulbecco’s Modified Eagle’s Medium (DMEM, 30-2214) supplemented with 10% Fetal Bovine Serum (FBS), and fed every other day. For electroporation, 5 x 10^5^ cells were used. For infections, 2.5 x 10^5^ cells were plated on a single well of a 6-well plate and infected the next day.

### Generation of SCYL1^AIv4^ alleles in Hek293T cells (eSPLIT and iSWAP)

eSPLIT and iSWAP alleles were engineered using CRISPR-Cas9 technology essentially as described previously[18]. For eSPLIT designs, plasmids encoding Cas9-P2A-eGFP under the control of the chicken β-actin (Cbh) promoter and sgRNAs specific for each locus under the control of the U6 promoter (px458 plasmid, Addgene #48138[2], **Additional Table 7**) were engineered and co-electroporated with single stranded DNA oligonucleotides as homology directed repair (HDR) templates into Hek293T cells using Amaxa 4D-Nucleofector X unit (Lonza). SF buffer and CM130 electroporation parameters were used as described by the manufacturer. For iSWAP alleles, Hek293T cells containing a single copy of the *SCYL1* gene *(SCYL1^+/LD^*, see below for details about the engineering of this cell line) were electroporated with PX458 and PX458-mCherry where the GFP cDNA was replaced by the cDNA encoding mCherry using conventional molecular biology techniques, each encoding sgRNA allowing for the removal and swap of intronic sequences. Guide sequences and HDR templates are described in **Additional Table 1**. 48 hours after electroporation, cells were single cell sorted based on fluorescence into 96 well plates (GFP for eSPLIT or GFP and mCherry for iSWAP). Clones were grown for 14-21 days, and replica plated for genomic DNA (gDNA) extraction and freeze down. Clones were characterized at the genomic level using PCR amplification and Sanger sequencing. Banding patterns (fragment sizes) of wildtype or properly targeted alleles are presented in **Additional Table 2**. The list of all models engineered in this manuscript are presented in **Additional Table 6**.

### Generation of SCYL1^+/LD^ Hek293T cells

Hek293T cells containing a single *SCYL1* gene was engineered by co-electroporation of 2 sgRNAs flanking the *SCYL1* gene (**Additional Fig. 5a – c**). PCR genotyping and Sanger sequencing confirmed the engineering of the cell line. A clone with a single copy of the *SCYL1* gene was isolated and used for subsequent studies using iSWAP designs.

### Hek293T-SCYL1-KO cell line

Hek293T cells deficient for SCYL1 have been described previously[19].

### Genomic DNA (gDNA) preparation from cultured cells

Cultured cells were washed three times with ice cold Dulbecco’s Phosphate Buffered Saline (DPBS). Washed cells were lysed with cell lysis buffer (NaCl, 10 mM; Tris HCL pH 7.5, 10mM; EDTA, 10 mM; 0.5% (w/v) Sodium Lauroyl Sarcosinate; Proteinase K, 500 µg/mL) for 16 h at 55°C. Lysed cells were incubated at 95°C for 10 minutes to heat inactivate Proteinase K.

### gDNA preparation from mouse toe or tail biopsies

Toe or tail biopsies were incubated in tail lysis buffer (KCl, 500 mM; Tris pH 8.3, 100 mM; Gelatin, 0.1 mg/mL; NP40, 1% (v/v); Tween-20, 1% (v/v); Proteinase K, 500 µg/mL) for 16 h at 55°C. Digested biopsies were incubated at 95°C for 10 minutes to heat inactivate Proteinase K.

### Generation of mouse models

Mouse models were engineered using CRISPR-Cas9 technology. Briefly, C57BL/6J zygotes were electroporated using the NEPA21 system (1 mm chamber) with Cas9 protein (100 ng/μL), crRNAs specific for each target sites (100 ng/μl), tracrRNAs (100 ng/μl) and HDR templates in the form of a ssDNA molecules (100 ng/μL). The following electroporation parameters were used: voltage, 40V; pulse length, 3 msec; pulse interval, 50 msec; number of pulses, 5; decay rate, 40%; and polarity +/-. Following electroporation, zygotes were cultured in KSOM medium for 1-2 hours and transferred to pseudo pregnant CD1 females. Pups obtained from these microinjections were characterized by PCR-based genotyping and Sanger sequencing using primers presented in **Additional Table 1**. Banding pattern obtained for each allele are presented in **Additional Table 2**. Sanger sequencing was performed using both forward and reverse primers.

### PCR-based genotyping

All PCR genotyping reactions were prepared on ice using Qiagen Taq polymerase (**Additional Table 5**), following the manufacturer’s recommendations. Thermal cycling was performed using the following protocol: 1) Initial denaturation at 95 °C for 10 minutes (1 cycle); 2) Denaturation at 95 °C for 30 seconds, annealing at 60 °C for 30 seconds, and extension at 72 °C for 2 minutes (35 cycles); and 3) Final extension at 72 °C for 10 minutes (1 cycle). For SCYL1^i6AIv4^ cell line model, PCR genotyping reactions were set up using Long Taq DNA polymerase (New England Biolabs), following the manufacturer’s recommendations. Thermal cycling was performed using the following protocol: 1) Initial denaturation at 95 °C for 10 minutes (1 cycle); 2) Denaturation at 95 °C for 30 seconds, annealing at 60 °C for 30 seconds, and extension at 72 °C for 7 minutes (38 cycles); and 3) Final extension at 72 °C for 10 minutes (1 cycle).

### Sanger sequencing

Sanger sequencing was performed as previously described [5, 20]. PCR products were treated with ExoSAP-IT™ PCR Product Cleanup Reagent (ThermoFisher Scientific) to remove excess primers and dNTPs. Sequencing reactions were prepared using the BigDye™ Terminator v3.1 Cycle Sequencing Kit (ThermoFisher Scientific) and run under standard thermal cycling conditions recommended by the manufacturer. Following cycle sequencing, reactions were purified using the BigDye™ XTerminator™ Purification Kit (ThermoFisher Scientific), which includes the XTerminator™ Solution and SAM™ Solution. Purified products were then loaded onto a SeqStudio™ Genetic Analyzer (ThermoFisher Scientific) for fragment analysis and detection. Sequencing data was analyzed using SnapGene® software (Dotmatics).

### Lentivirus production and infection

Lentivirus expressing Cre recombinase was produced by co-transfecting HEK293T cells with plasmids encoding the Cre recombinase and mCherry under the control of CMV promoter (pLenti-mCherry-Cre-blast (Addgene#179390)) together with psPAX2 (Addgene #12260) and pMD2.G (addgene #12259) which encode Gag, Pol, Rev proteins for lentiviral packaging and VSV-G envelope respectively. 48 hours after transfection, supernatants were collected and filtered through a 0.45 μm syringe filter. Freshly made viruses were used within 48 hours to infect engineered Hek293T cells. Cells were infected with an equal amount (2 mL) of filtered supernatant containing the viral particles and polybrene (8 μg/mL). 72 hours post-infection, media was changed for medium containing blasticidin (10 µg/mL) where indicated. 6 days following infection, cells were washed twice with DPBS and collected for gDNA preparation or lysed for western blotting.

### Western blotting

Cells or tissues were lysed in NP-40 lysis buffer (Triton X-100, Tris-HCL, NaCl, PhosStop, Roche Complete protease inhibitor tablets) for 30 min on ice. Tissues were further homogenized using a tissue homogenizer, on ice for 30 seconds. Cell or tissue lysates were cleared by centrifugation and protein content assessed using BCA assay (Pierce). 5 µg of total proteins were loaded on 4-12% or 10% Criterion™ XT Bis-Tris Protein Gels and transferred to nitrocellulose membranes. Membranes were incubated in Tris Buffered Saline with Tween 20 (TBST) 1X (prepared from TBST-10X Cell signaling technologies, Cat#9997) containing 5% nonfat dry milk for 60 minutes. Membranes were subsequently incubated with primary antibodies, overnight, 4°C. Following incubation, membranes were washed 3 times for 5 minutes each with TBST. Following the third wash, membranes were incubated with HRP conjugated secondary antibodies in TBST 1X containing 5% nonfat dry milk for 60 minutes at 20°C. Membranes were then washed 3 times in TBST 1x and proteins detected using ECL detection assay from Bio-Rad.

### Mouse husbandry

Mice were housed in an Association for Assessment and Accreditation of Laboratory Animal Care accredited facility and maintained in accordance with the National Institutes of Health Guide for the Care and Use of Laboratory Animals. Animal experiments were reviewed and approved by the Indiana University Institutional Animal Care and Use Committee.

### Mouse breeding scheme

Genetically engineered mouse models were outbred for two generations to C57BL/6J mice (Jackson Laboratories) prior to crossing them to Cre or FLP driver to generate mice expressing the Cre or the FLP transgene (Cre+ or FLP+) and heterozygous for the conditional allele (+/AIv4 or +/AIv5). In parallel, mice homozygous for the conditional alleles were generated by heterozygous intercrosses. These mice were then bred to Cre+;+/AIv4Δ or FLP+;AIv5Δ mice to generate mice homozygous for the recombined allele. Of note, mosaic deletion in Cre expressing mice was observed in several animals, resulting in partial, but significant inactivation of the engineered alleles, which is likely the result of cellular mosaicism of Cre expression rather than the recombination inefficiency of the AIv4 cassette as detailed previously[10].

## ACKNOWLEDGEMENT

The authors thank Judy Hallet from the Purdue University Transgenic Facility for the generation of Rgma founder animals. This research was made possible by an award from the Indiana University School of Medicine and the Department of Medical and Molecular Genetics at Indiana University School of Medicine. Plasmid pLenti_mCherry_Cre_Blast was a gift from Moshe Szyf (Addgene plasmid #179390)[1]. pMD2.G and psPAX2 (Addgene # 12259 and 12260) were a gift from Didier Trono via Addgene. PX458 (Addgene # 48138) was a gift from Feng Zhang[2].

## Additional Figures

**Additional figure 1. Genomic characterization of engineered cell line models.**

Sanger sequencing traces illustrating properly targeted eSPLIT clones. All correctly targeted homozygous clones (AIv4/AIv4) were validated with Sanger Sequencing.

**Additional Figure 2. Cre-mediated recombination of functional e-SPLIT AIv4 alleles.**

a) Cre-mediated recombination was performed using lentiviral delivery of Cre recombinase using pLenti_mCherry_Cre-Blast.

b) Representative pictures of HEK293T cells 72 hours after infection. Cells with red fluorescence showed successfully infected cells.

c) Sanger sequencing traces of recombined HDR alleles following lentiviral delivery of Cre recombinase in e4_HDRb_1C11^AIv4Δ/AIv4Δ^, e4_HDRc_C6^AIv4Δ/AIv4Δ^, e6_HDRa_1E6^AIv4Δ/AIv4Δ^ and e16_HDRa_1E6^AIv4Δ/AIv4Δ^ clones.

**Additional figure 3. Sanger sequencing of the Exon4^AIv4^ clones in three frames.**

a) Sanger sequencing traces of all clones used in Fig. 3, testing the impact of inserting the AIv4 cassette within all three frames: e4_HDRb_1C11^AIv4/AIv4^, e4_HDRd_3C3^AIv4/AIv4^ and e4_HDRe_1A12 ^AIv4/AIv4^.

b) Sanger sequencing traces of Cre recombined alleles in e4_HDRb_1C11^AIv4Δ/AIv4Δ^, e4_HDRd_3C3^AIv4Δ/AIv4Δ^ and e4_HDRe_1A12 ^AIv4Δ/AIv4Δ^ clones.

**Additional figure 4. Rufy1^AIv4^ mouse model design strategy and validation at the genomic level using Sanger sequencing.**

a) Schematic representation of the *Rufy1* gene and generation of the *Rufy1^AIv4-1^*, *Rufy1^AIv4-2^* and Rufy1^Del76^ alleles. Guide cut sites, Rufy1_AIv4-1_G01 and Rufy1_AIv4-2_G02 are indicated by downward black arrows. Two black arrows mark the 76-nucleotide deletion at the Intron 1–Exon 2 junction of the *Rufy1* gene. Blue arrow indicates location of forward genotyping primer. Green arrow indicates location of reverse genotyping primer.

b) Correctly targeted allele for *Rufy1*^AIv4-1^ was validated using Sanger Sequencing.

c) Correctly targeted allele for *Rufy1*^AIv4-2^ was validated with Sanger Sequencing.

d) Sanger sequencing traces of *Rufy1*^Del76/Del76^ locus.

e) Recombined allele for *Rufy1_AIv4-2^AIv4Δ/AIv4Δ^* after Cre mediated recombination was validated with Sanger Sequencing.

f) Schematic representation of the *Rgma* gene and generation of the Rgma^AIv4/AIv4^ allele. Guide cut site, *Rgma*^AIv4^_G01 is indicated by downward black arrows. Blue arrow indicates location of forward genotyping primer, *Rgma*^AIv4^_F51. Green arrow indicates location of reverse genotyping primer, *Rgma*^AIv4^_R52.

g) Sanger sequencing traces of the *Rgma*^AIv4/AIv4^ locus.

h) Sanger sequencing traces of the *Rgma^AIv4Δ/AIv4Δ^ locus following* Cre-mediated recombination.

**Additional figure 6. Engineering of Hek293T cells with a single SCYL1 allele and details iSWAP design strategies.**

A) Schematic representation of the *SCYL1* gene and generation of the SCYL1_LD allele.

B) Validation of Hek293T cells clones bearing a single functional allele of *SCYL1*. Genotyping (left) and Western Blot (right) illustrating the proper modification of the locus and expression of SCYL1 protein compared to parental cell line.

C) Schematic representation of the *SCYL1* gene and generation of *SCYL1^i2AIv4^* allele. Two guide cut sites, hSCYL1_i2_AIv4_G1 (i2_G1) and hSCYL1_i2_AIv4_G2 (i2_G2) are indicated by downward black arrows. Blue arrow indicates location of forward genotyping primer. Green arrow indicates location of reverse genotyping primer.

D) Schematic representation of the *SCYL1* gene and generation of *SCYL1^i3AIv4^*allele. Two guide cut sites, hSCYL1_i3_AIv4_G1 (i3_G1) and hSCYL1_i3_AIv4_G2 (i3_G2) are indicated by downward black arrows. Blue arrow indicates location of forward genotyping primer. Green arrow indicates location of reverse genotyping primer.

E) Schematic representation of the *SCYL1* gene and generation of *SCYL1^i6AIv4^*allele. Two guide cut sites, hSCYL1_i6_AIv4_G1 (i6_G1) and hSCYL1_i6_AIv4_G2 (i6_G2) are indicated by downward black arrows. Blue arrow indicates location of forward genotyping primer. Green arrow indicates location of reverse genotyping primer.

**Additional figure 7. Design strategy and Sanger Sequencing for Nipsnap1^i2AIv4^ mouse model.**

A) Schematic representation of the *Nipsnap1* gene and generation of the Nipsnap1^i2AIv4^ allele. Two guide cut sites, *Nipsnap1*^i2AIv4^_G01 and *Nipsnap1*^i2AIv4^_G02 are indicated by downward black arrows. Blue arrow indicates location of forward genotyping primer, *Nipsnap1*^i2AIv4^_F51. Green arrow indicates location of reverse genotyping primer, *Nipsnap1*^i2AIv4^_R52.

B) Correctly targeted allele for *Nipsnap1*^i2AIv4/AIv4^ was validated with Sanger Sequencing.

C) Recombined allele for *Nipsnap1*^i2*AIv4Δ/AIv4Δ*^ after Cre mediated recombination was validated with Sanger Sequencing.

**Additional figure 8. Design strategy and Sanger Sequencing for Stmn2^AIv5^ mouse model.**

A) Schematic representation of the *Stmn2* gene and generation of the *Stmn2*^AIv5^ allele. Guide cut site, *Stmn2*^AIv5^_G01 is indicated by downward black arrow. Blue arrow indicates location of forward genotyping primer, *Stmn2*^AIv5^_F53. Green arrow indicates location of reverse genotyping primer, *Stmn2*^AIv5^_R54.

B) Correctly targeted allele for *Stmn2*^AIv5/AIv5^ was validated with Sanger Sequencing.

C) Recombined allele for *Stmn2^AIv5Δ/AIv5Δ^* after FLPO mediated recombination was validated with Sanger Sequencing.

## Notes

### Competing Interest Statement

The authors have declared no competing interest.

## REFERENCES

1. Sapozhnikov DM, Szyf M: Unraveling the functional role of DNA demethylation at specific promoters by targeted steric blockage of DNA methyltransferase with CRISPR/dCas9. Nat Commun 2021, 12(1):5711.

2. Ran FA, Hsu PD, Wright J, Agarwala V, Scott DA, Zhang F: Genome engineering using the CRISPR-Cas9 system. Nat Protoc 2013, 8(11):2281–2308.

3. Erhardt V, Hartig E, Lorenzo K, Megathlin HR, Tarchini B, Hosur V: Systematic optimization and prediction of cre recombinase for precise genome editing in mice. Genome Biol 2025, 26(1):85.

4. Cassidy A, Pelletier S: CRISPR-Cas9-mediated insertion of a short artificial intron for the generation of conditional alleles in mice. STAR Protoc 2023, 4(1):102116.

5. Cassidy AM, Thomas DB, Kuliyev E, Chen H, Pelletier S: One-step generation of a conditional allele in mice using a short artificial intron. Heliyon 2022, 8(12):e12630.

6. Wu SS, Lee H, Szep-Bakonyi R, Colozza G, Boese A, Gert KR, Hallay N, Lee JH, Kim J, Zhu Y et al: SCON-a Short Conditional intrON for conditional knockout with one-step zygote injection. Exp Mol Med 2022, 54(12):2188–2199.

7. Guzzardo PM, Rashkova C, Dos Santos RL, Tehrani R, Collin P, Burckstummer T: A small cassette enables conditional gene inactivation by CRISPR/Cas9. Scientific reports 2017, 7(1):16770.

8. Pelletier S, Gingras S, Howell S, Vogel P, Ihle JN: An early onset progressive motor neuron disorder in Scyl1-deficient mice is associated with mislocalization of TDP-43. The Journal of neuroscience : the official journal of the Society for Neuroscience 2012, 32(47):16560–16573.

9. Economides AN, Frendewey D, Yang P, Dominguez MG, Dore AT, Lobov IB, Persaud T, Rojas J, McClain J, Lengyel P et al: Conditionals by inversion provide a universal method for the generation of conditional alleles. Proc Natl Acad Sci U S A 2013, 110(34):E3179–3188.

10. Schwenk F, Baron U, Rajewsky K: A cre-transgenic mouse strain for the ubiquitous deletion of loxP-flanked gene segments including deletion in germ cells. Nucleic Acids Res 1995, 23(24):5080–5081.

11. Wu Y, Wang C, Sun H, LeRoith D, Yakar S: High-efficient FLPo deleter mice in C57BL/6J background. PloS one 2009, 4(11):e8054.

12. Cassidy A, Onal M, Pelletier S: Novel methods for the generation of genetically engineered animal models. Bone 2022, 167:116612.

13. Gurumurthy CB, O’Brien AR, Quadros RM, Adams J, Jr., Alcaide P, Ayabe S, Ballard J, Batra SK, Beauchamp MC, Becker KA et al: Reproducibility of CRISPR-Cas9 methods for generation of conditional mouse alleles: a multi-center evaluation. Genome Biol 2019, 20(1):171.

14. Lanza DG, Gaspero A, Lorenzo I, Liao L, Zheng P, Wang Y, Deng Y, Cheng C, Zhang C, Seavitt JR et al: Comparative analysis of single-stranded DNA donors to generate conditional null mouse alleles. BMC Biol 2018, 16(1):69.

15. Andersson-Rolf A, Mustata RC, Merenda A, Kim J, Perera S, Grego T, Andrews K, Tremble K, Silva JC, Fink J et al: One-step generation of conditional and reversible gene knockouts. Nat Methods 2017, 14(3):287–289.

16. Soulez M, Saba-El-Leil MK, Turgeon B, Mathien S, Coulombe P, Klinger S, Rousseau J, Levesque K, Meloche S: Reevaluation of the Role of Extracellular Signal-Regulated Kinase 3 in Perinatal Survival and Postnatal Growth Using New Genetically Engineered Mouse Models. Molecular and cellular biology 2019, 39(6).

17. Ohtsuka M, Sato M, Miura H, Takabayashi S, Matsuyama M, Koyano T, Arifin N, Nakamura S, Wada K, Gurumurthy CB: i-GONAD: a robust method for in situ germline genome engineering using CRISPR nucleases. Genome Biol 2018, 19(1):25.

18. Kuliyev E, Gingras S, Guy CS, Howell S, Vogel P, Pelletier S: Overlapping Role of SCYL1 and SCYL3 in Maintaining Motor Neuron Viability. The Journal of neuroscience : the official journal of the Society for Neuroscience 2018, 38(10):2615–2630.

19. Gingras S, Kuliyev E, Pelletier S: SCYL1 does not regulate REST expression and turnover. PloS one 2017, 12(6):e0178680.

20. Cassidy AM, Kuliyev E, Thomas DB, Chen H, Pelletier S: Dissecting protein function in vivo: Engineering allelic series in mice using CRISPR-Cas9 technology. Methods Enzymol 2022, 667:775–812.

