## Supplementary figures and images for "eSPLIT and iSWAP: CRISPR-Mediated Conditional Allele Engineering with Short Artificial Introns"

### Supplemental Figures

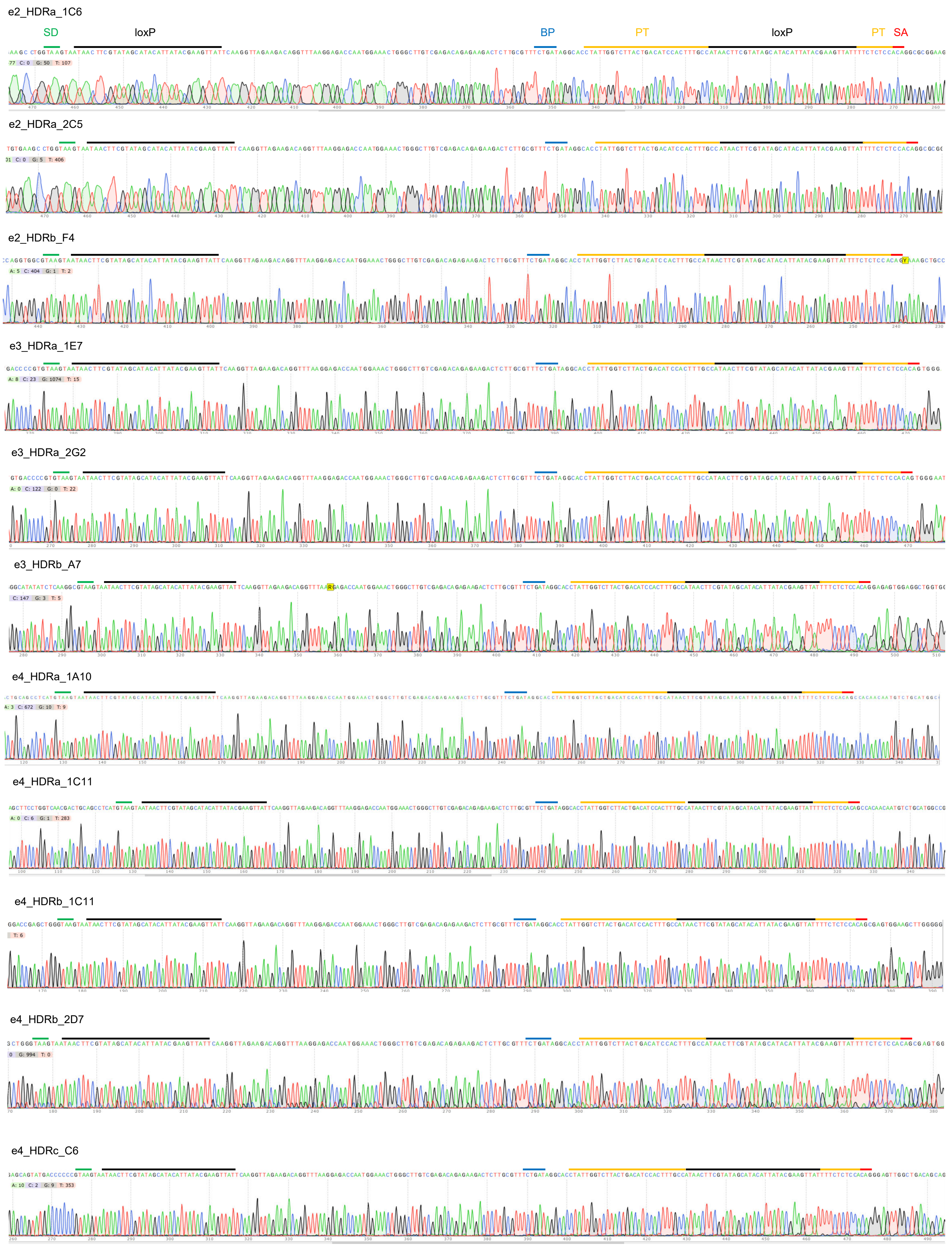

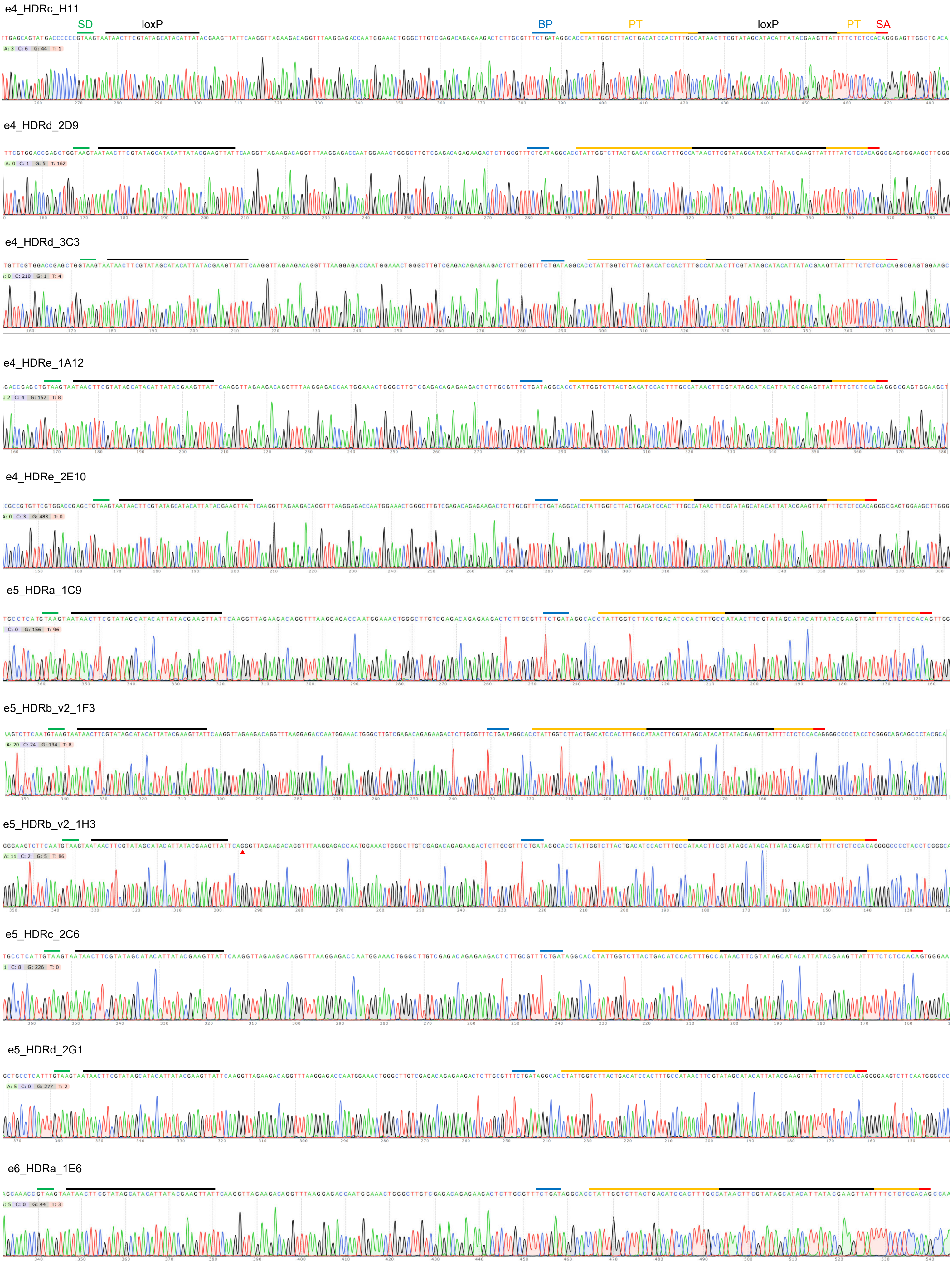

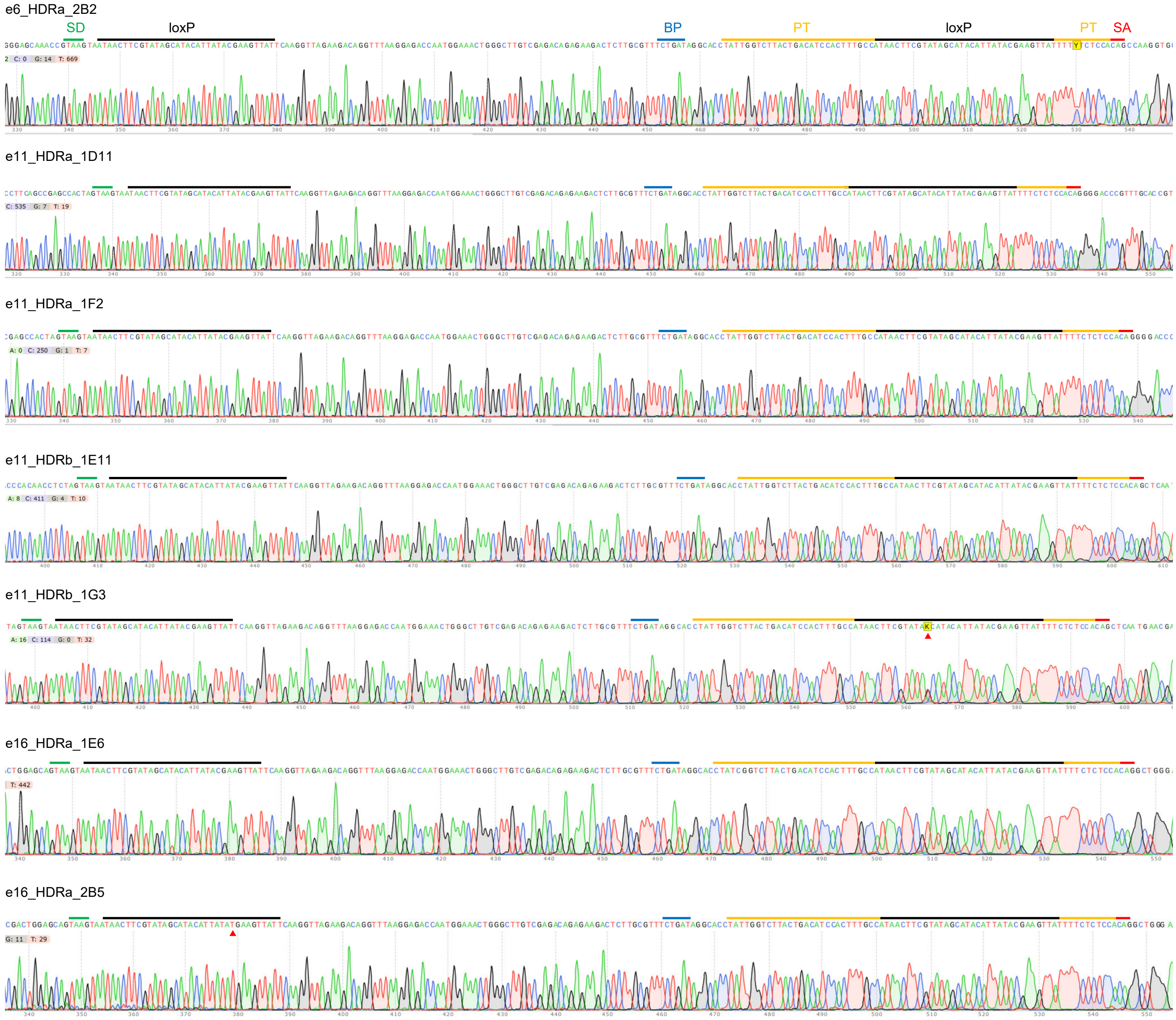

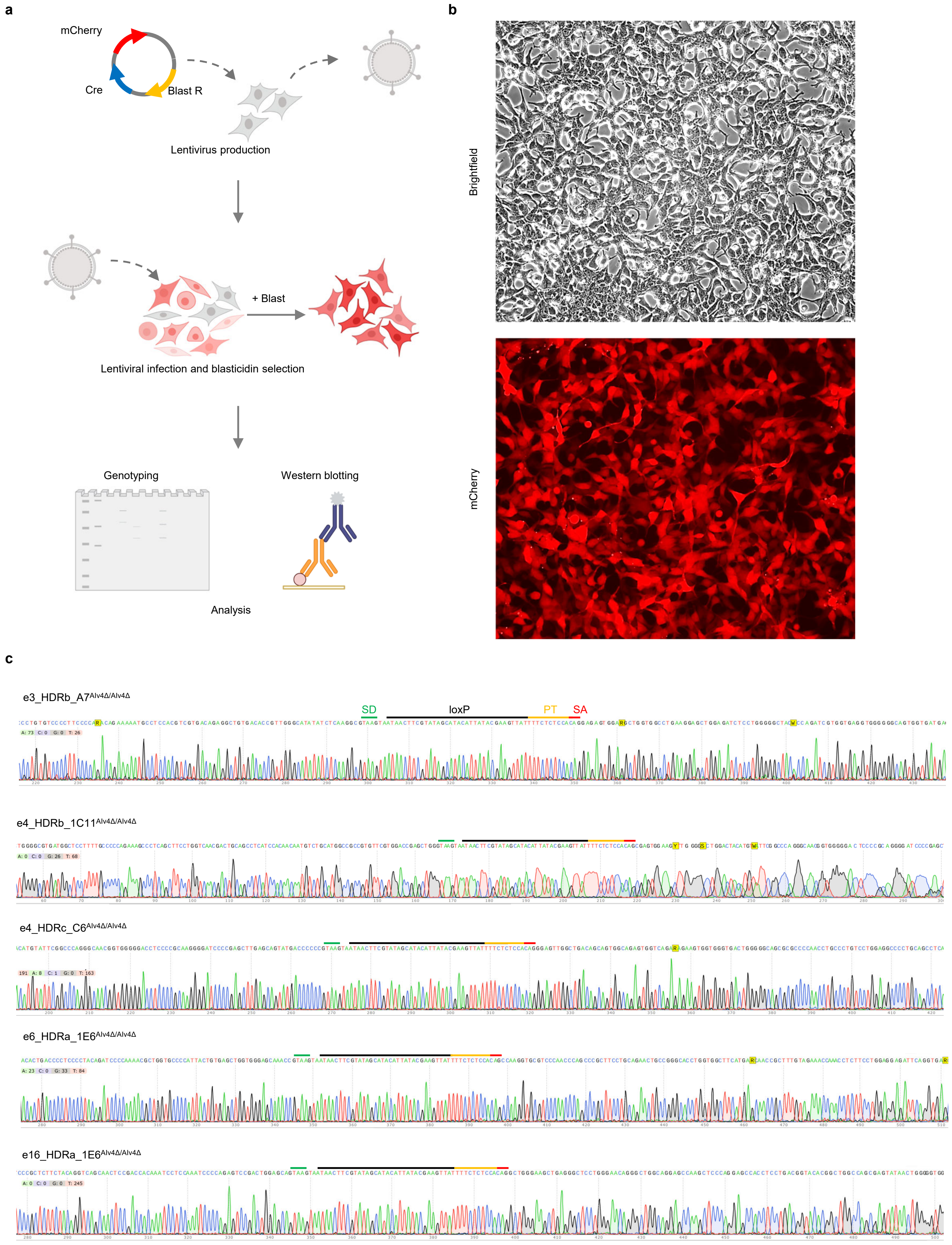

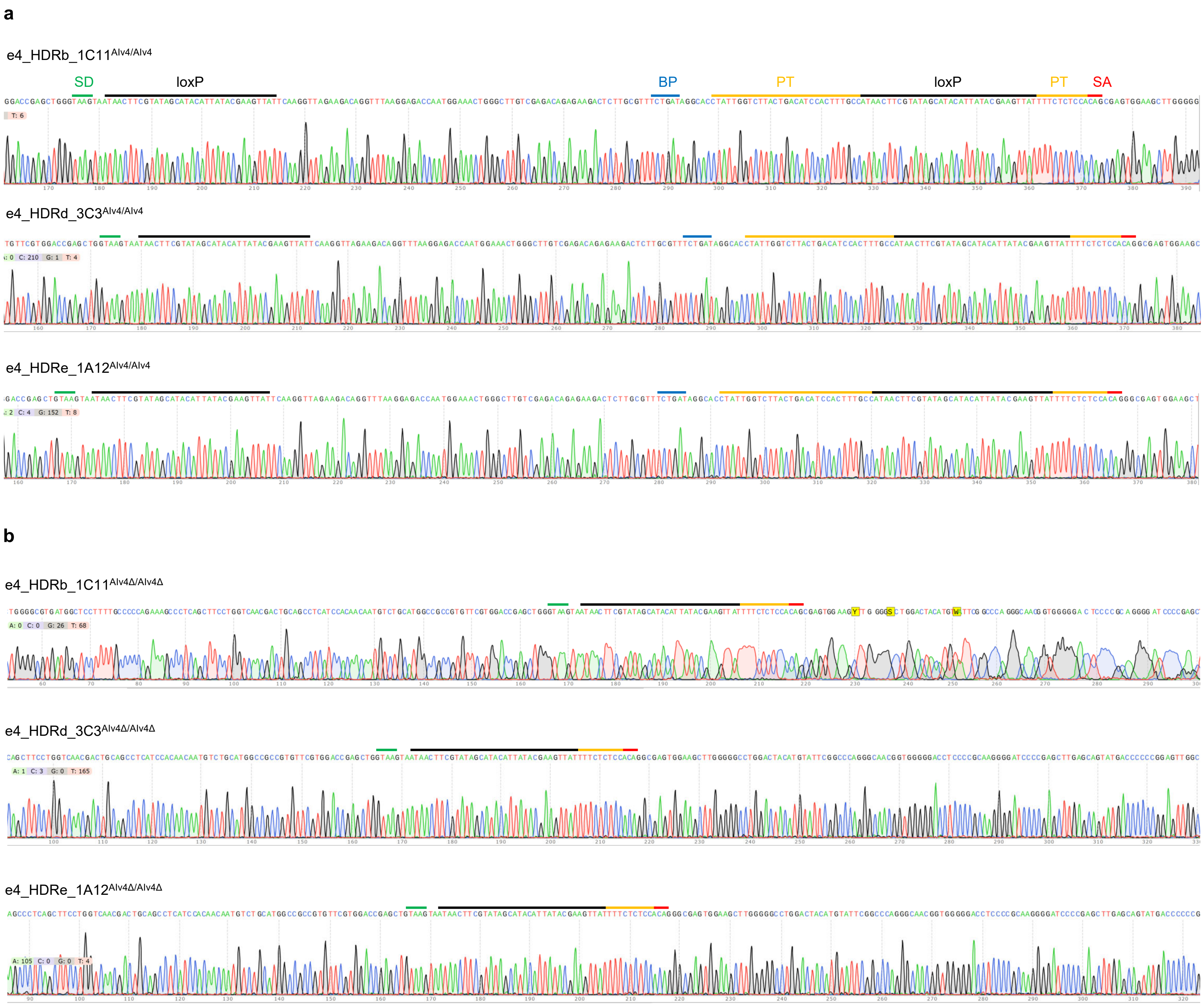

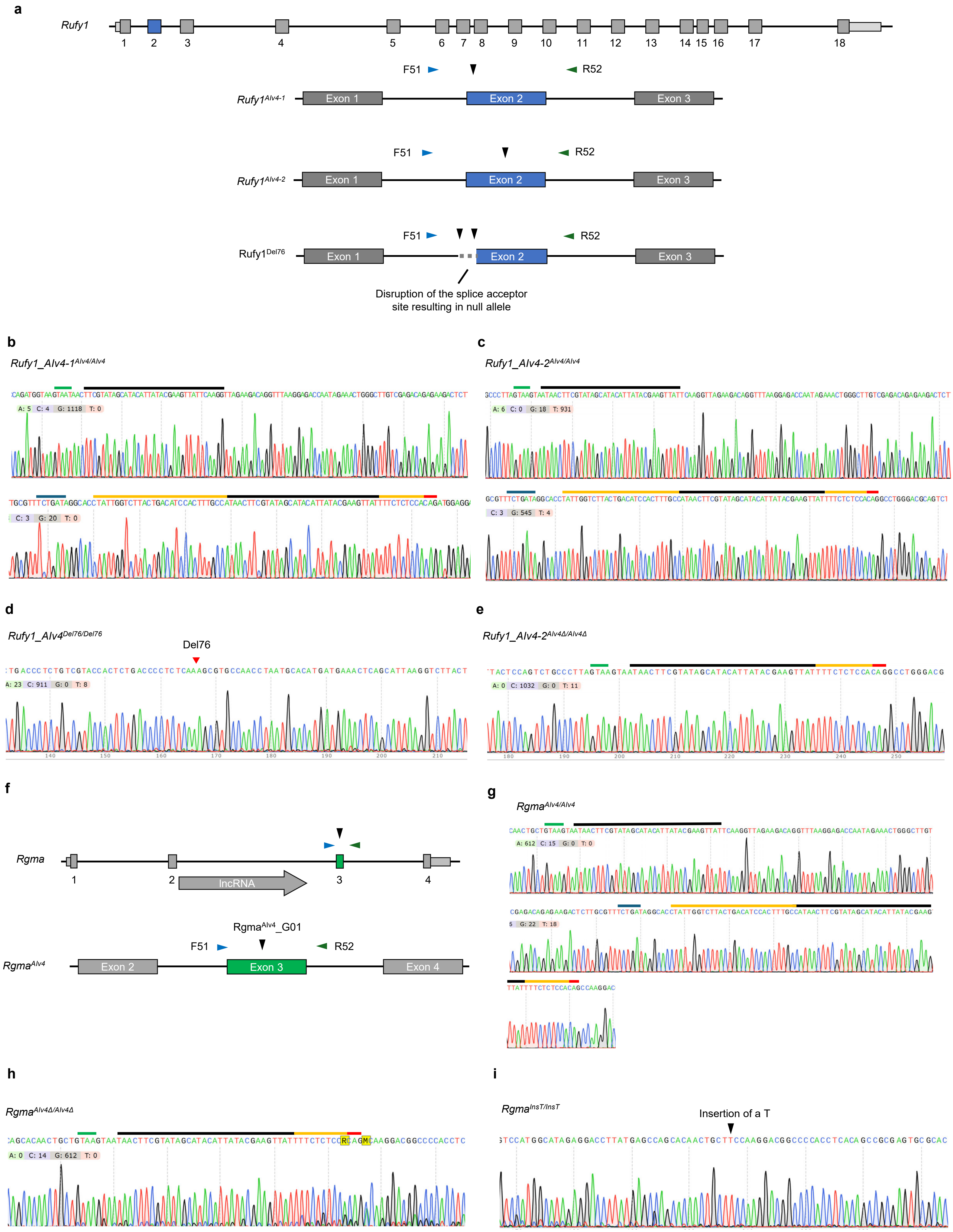

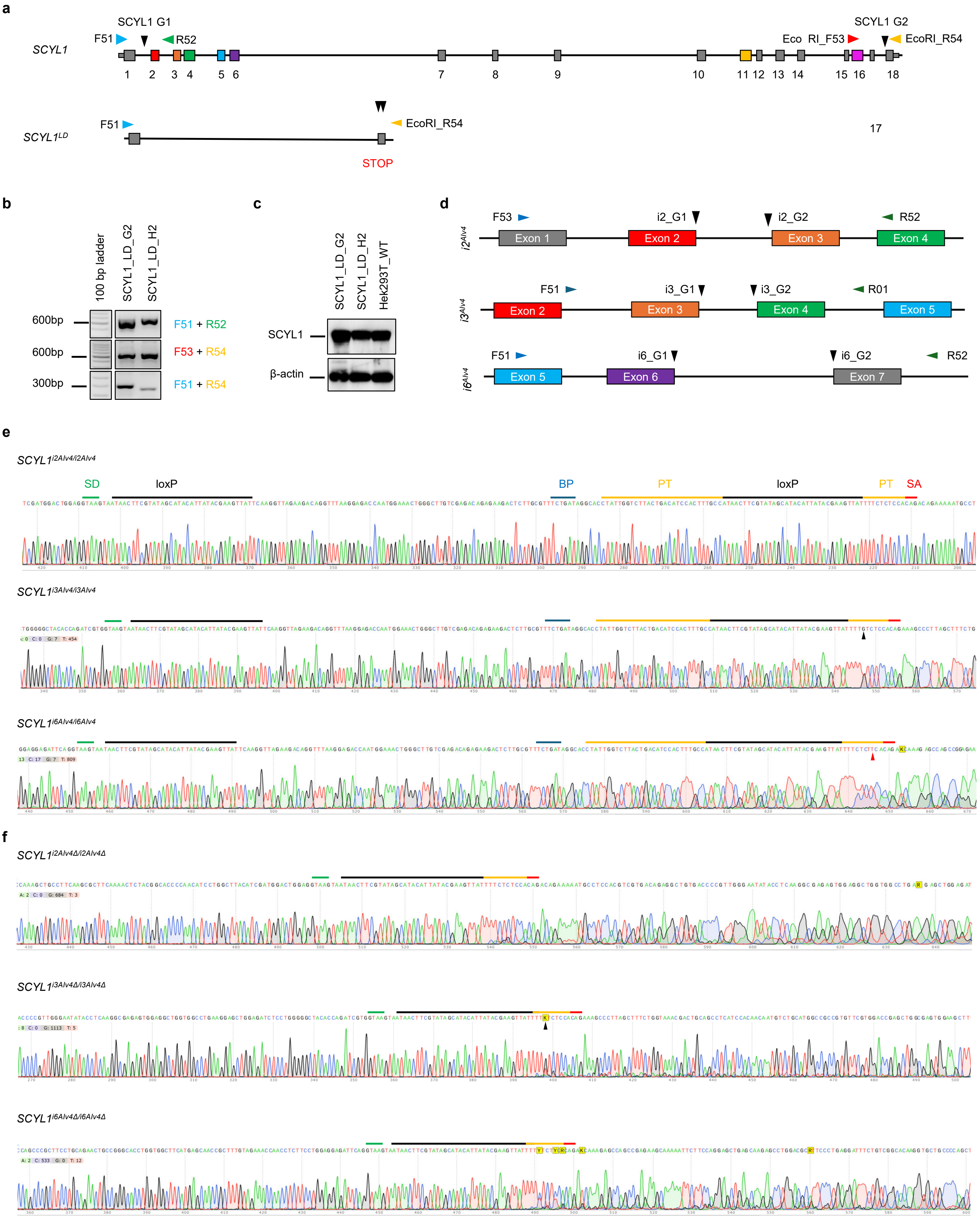

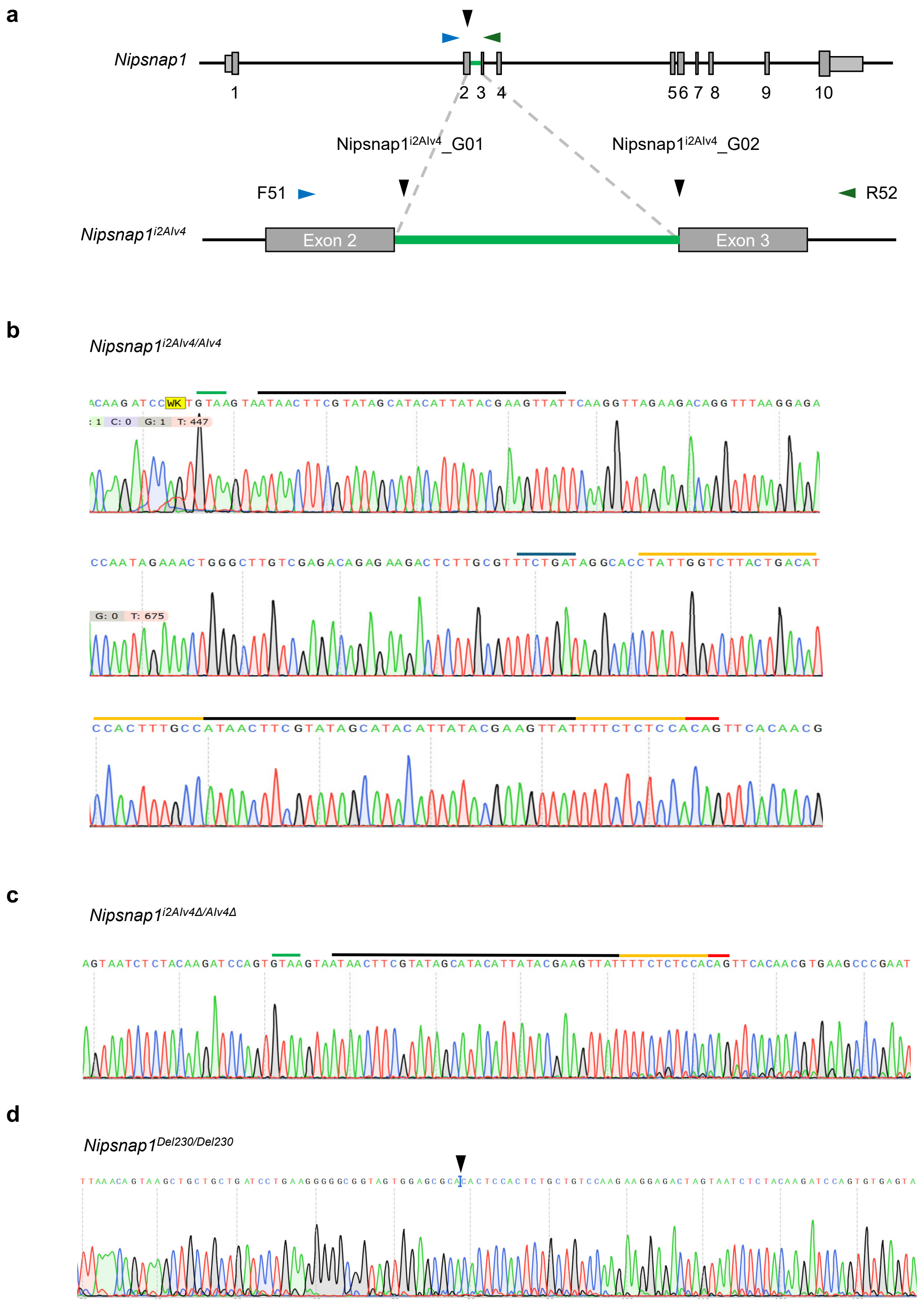

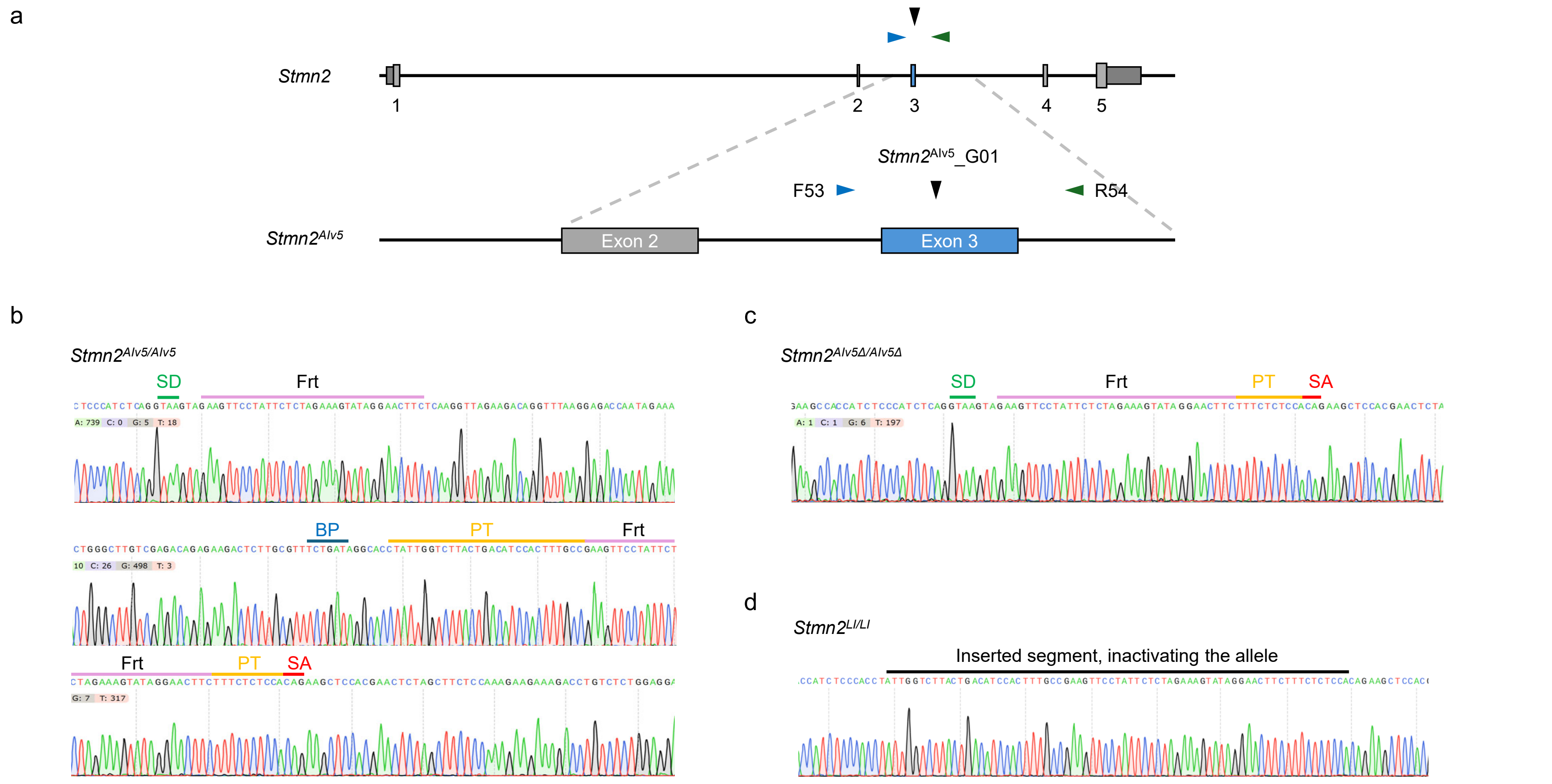
